## Supplementary figures and images for "Apical annuli are specialised sites of post-invasion secretion of dense granules in *Toxoplasma*"

### Figure S1

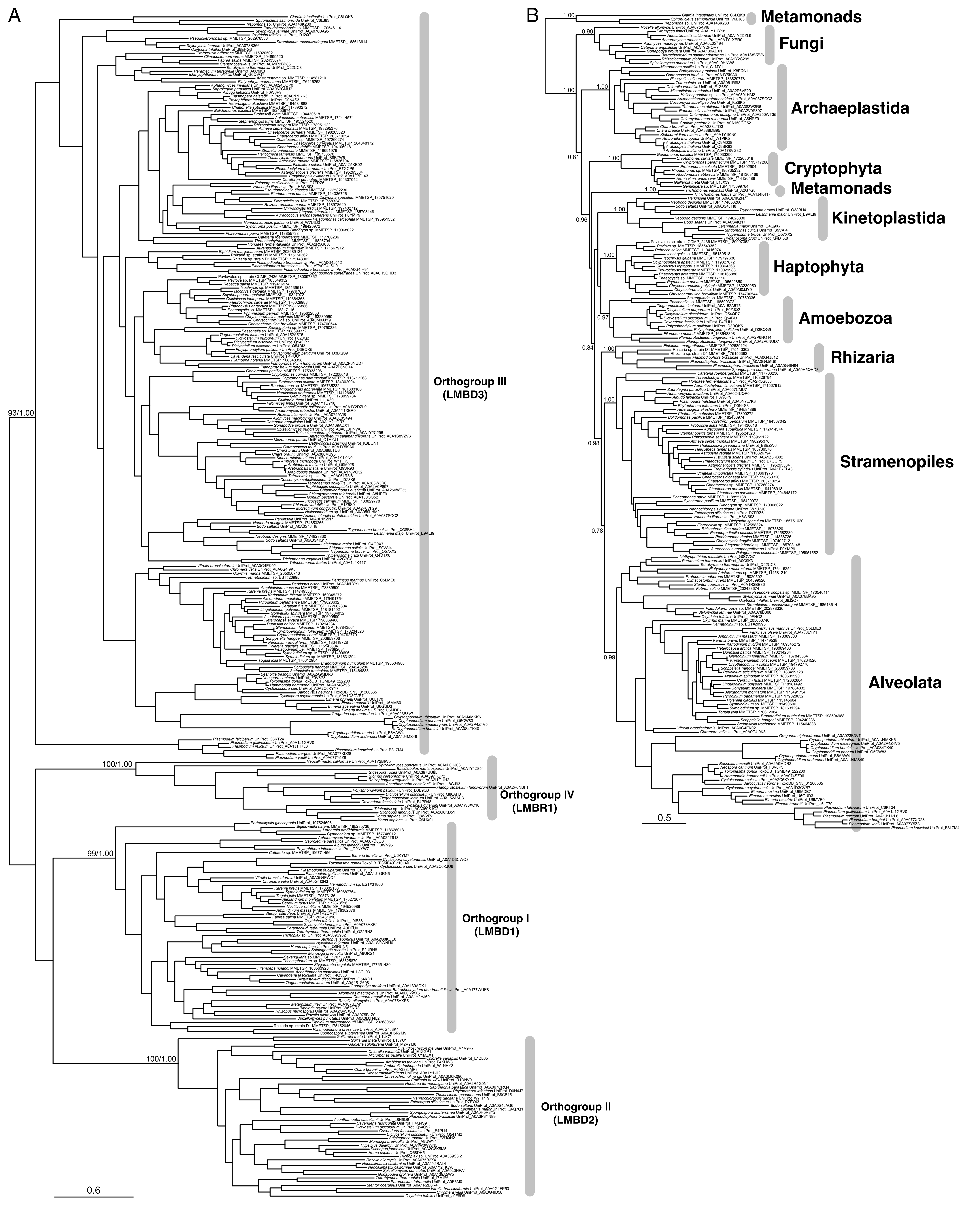

### Figure S2

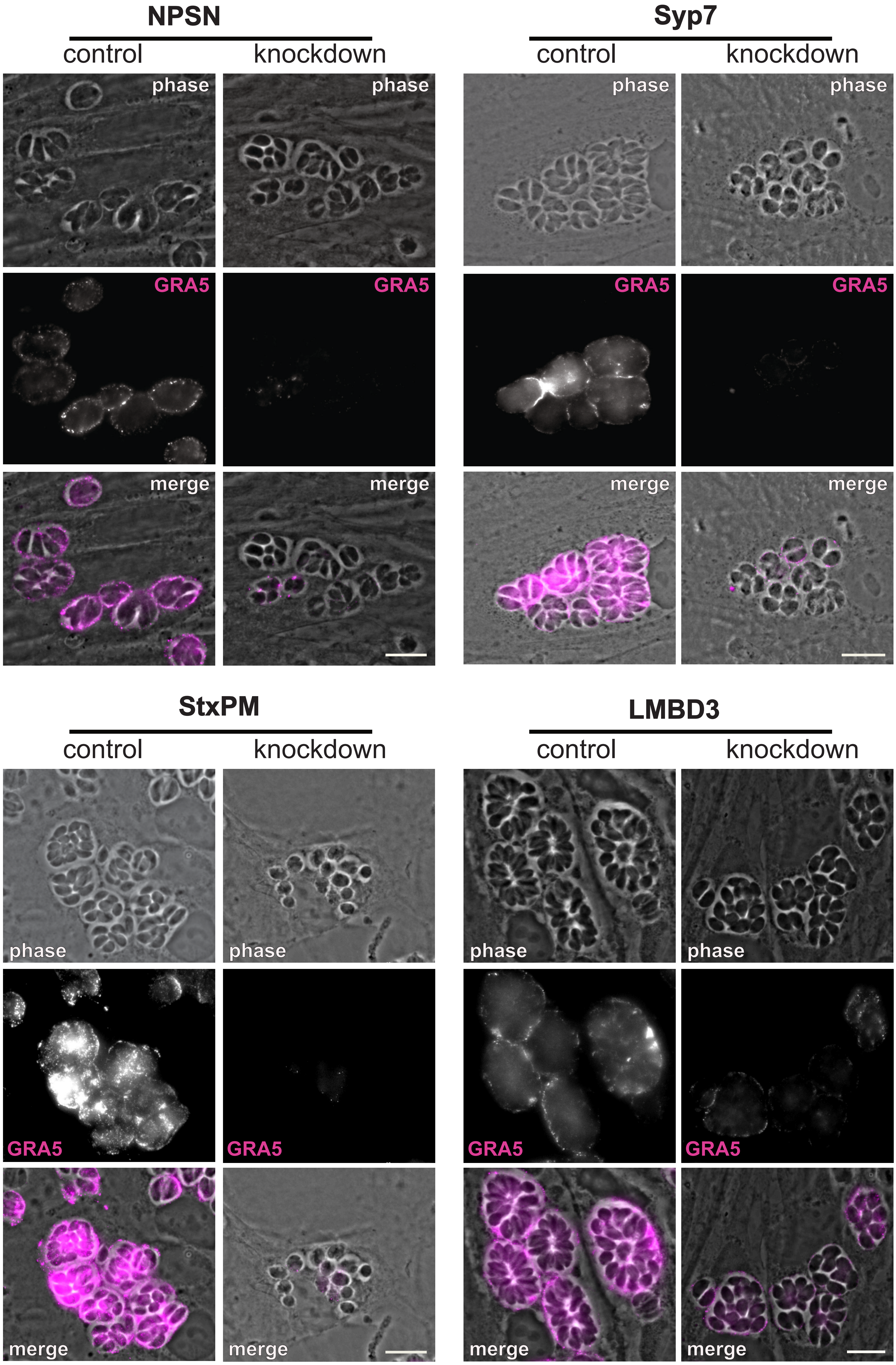

### Figure S3

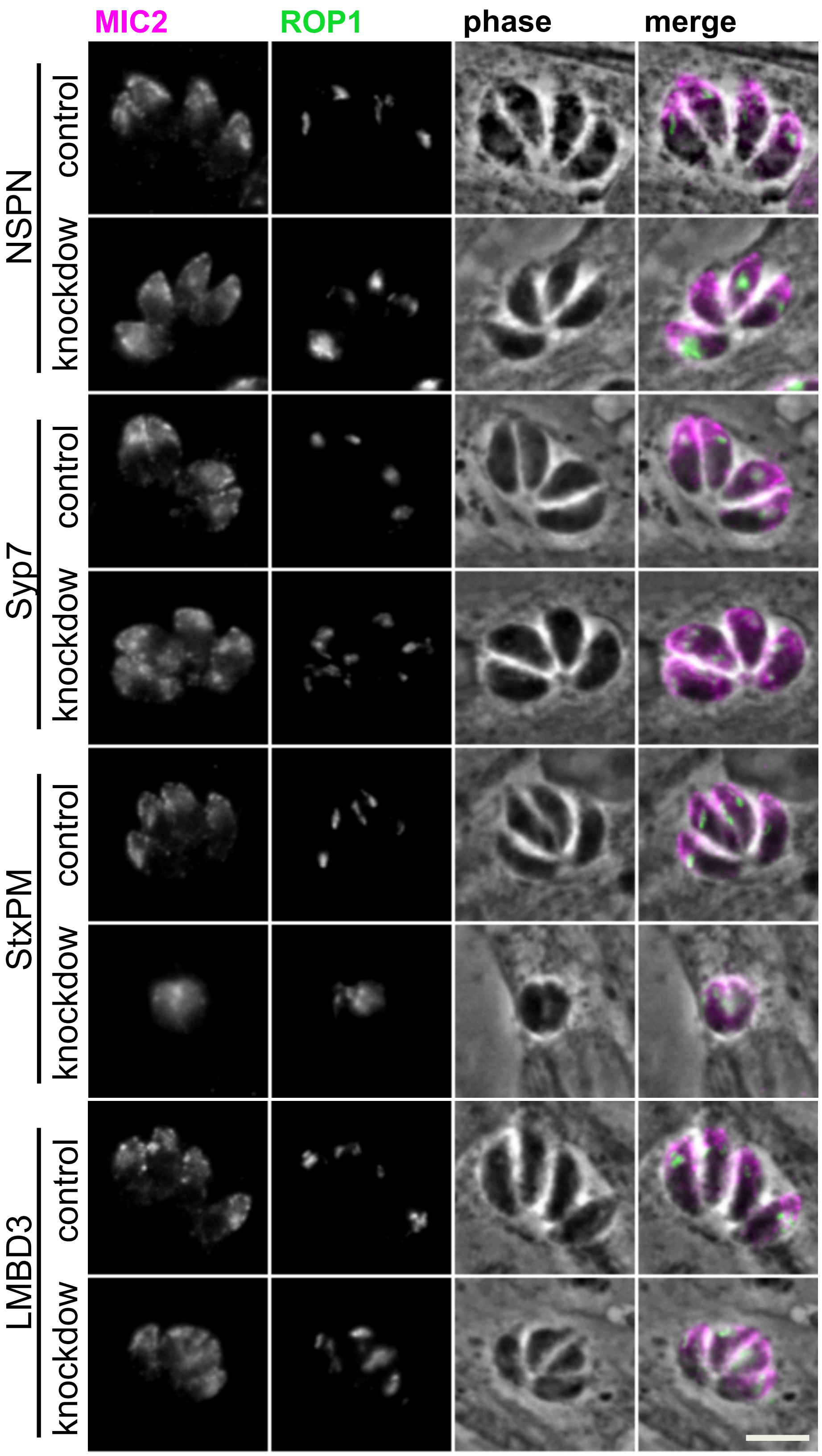

### Figure S4

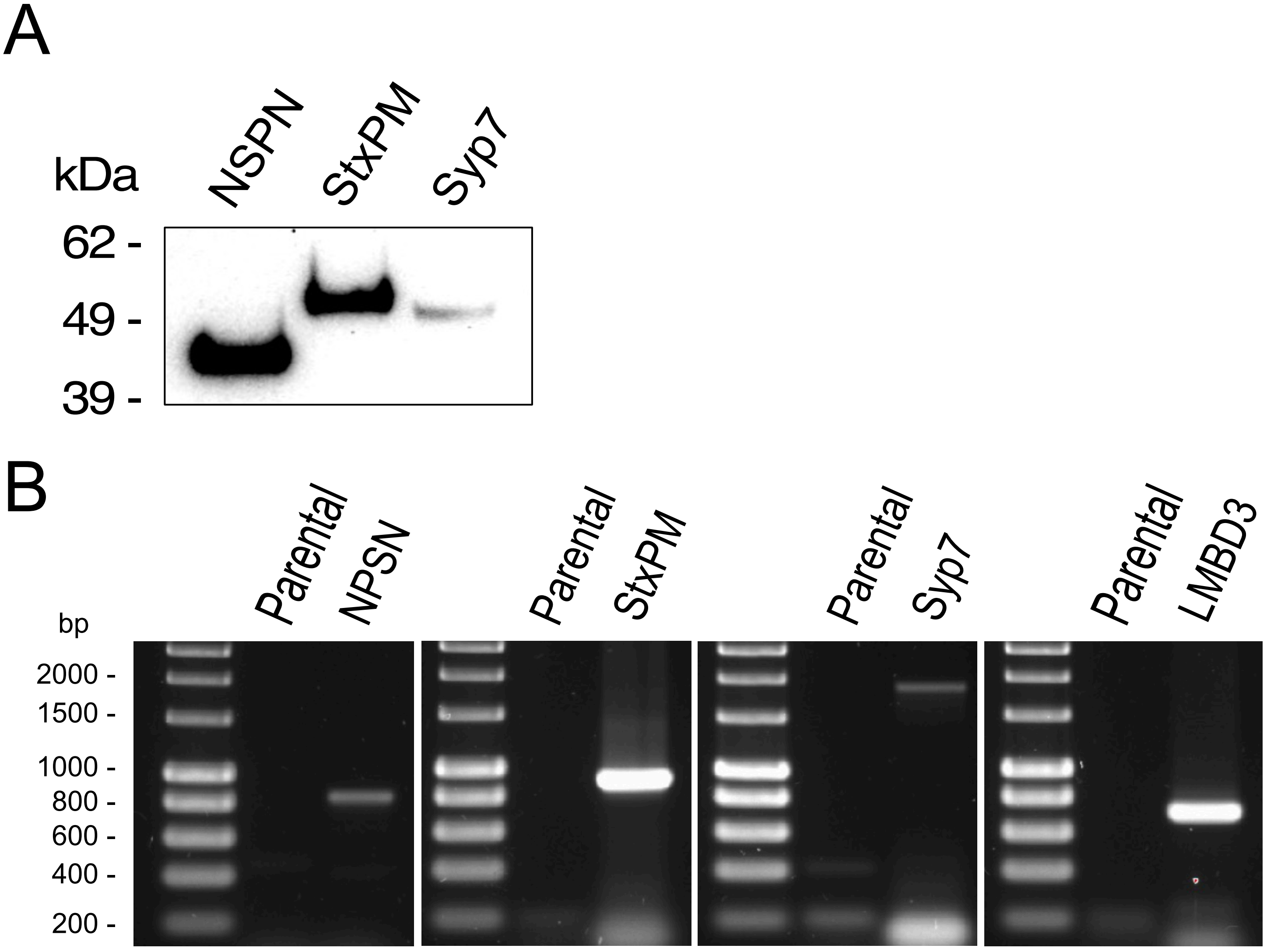
