## Supplementary material for "Apical annuli are specialised sites of post-invasion secretion of dense granules in *Toxoplasma*": Data file S1

### Toxo annuli quant prot

Thomas Krueger

2022-07-23

**Abstract**

This workflow describes the analysis of differential protein abundance in four different Toxoplasma gondii cell lines. Input data to this workflow is the PSM file generated by Proteome Discoverer 3.0, derived from SPS-MS3 analysis of TMT-labeled peptides (2 batches; 16plex each). The workflow covers initial quality analysis of MS data, PSM quality filtering, definition of individual datasets of interest, assessment and removal of missing data, feature aggregation from PSM to protein level, data normalisation across replicates and treatments,power analysis, linear model fitting, and generation of output files and plots with logfold changes and statistical signifance for all treatment contrasts of interest.

```
library(tidyverse)
library(scales)
library(cowplot)
library(QFeatures)
library(NormalyzerDE)
library(magrittr)
library(eulerr)
library(FactoMineR)
library(factoextra)
library(limma)
library(ggforce)
library(ggExtra)
library(pwr)
library(esc)
library(Biostrings)
library(writexl)
library(conflicted)

#command preference in case of package conflicts
conflicts_prefer(magrittr::set_names)
conflicts_prefer(dplyr::filter)
conflicts_prefer(matrixStats::colMedians)
conflicts_prefer(dplyr::rename)
conflicts_prefer(matrixStats::rowSds)
```

```
#loading psm level data
df_PSM <-
  read_delim(file = "input/P1218_Toxo_Annuli_PSMs.txt",
             delim = "\t",
             na = c("", "NA"),
             col_types = NULL) %>%
  data.frame()
```

```r
nm_batches <- c("Batch_1", "Batch_2")
nm_cell_lines <- c("TgLMBD3", "TgNPSN", "TgStxPM", "TgSyp7")

#list of experiments and contrasts of interest
contrasts <-
  read_csv(file = "input/Toxo_annuli_contrasts.csv",
           col_types = "c") %>%
  as.data.frame()

contrasts
```

```
##   experiment treat1  treat2
## 1    TgLMBD3     KD control
## 2     TgNPSN     KD control
## 3    TgStxPM     KD control
## 4     TgSyp7     KD control
```

```r
#shortening "Abundance" column names to reflect only TMT channel
df_PSM <-
  df_PSM %>%
  rename_with(.,
              ~ gsub(pattern = "Abundance.",
                     replacement = "",
                     x = .))

#Creating ExpSet ID column since data comes from two separate LCMS batch files
df_PSM$Spectrum.File %>% as.factor() %>% levels
```

```
## [1] "P1218_Toxo_Annuli_Test1.raw" "P1218_Toxo_Annuli_Test2.raw"
```

```r
df_PSM$ExpSet <- sub(".raw*", "", df_PSM$Spectrum.File) %>% as.factor()
df_PSM$ExpSet <- recode_factor(df_PSM$ExpSet,
                               P1218_Toxo_Annuli_Test1 = "Batch_1",
                               P1218_Toxo_Annuli_Test2 = "Batch_2")
df_PSM$ExpSet %>% levels()
```

```
## [1] "Batch_1" "Batch_2"
```

```r
#PSM quality thresholds
Isol_In <- 75
SN <- 10
SPS_Mass_match <- 70

#TMT channel order
TMT_order <- c("126", "127N", "127C", "128N",
               "128C", "129N", "129C", "130N",
               "130C", "131N", "131C", "132N",
               "132C", "133N", "133C", "134N")
```

```r
#Creating individual PSM datasets for each batch to check on the quality in both runs
ls_Batch_PSM <-
```

```r
  list(df_PSM %>% filter(ExpSet == "Batch_1"),
       df_PSM %>% filter(ExpSet == "Batch_2")) %>%
  set_names(nm_batches)


#overview of PSM quality
graphs_qual <-
  lapply(X = names(ls_Batch_PSM),
         FUN = function(i) {

           #get object
           obj <- ls_Batch_PSM[[i]]

           #time to accumulate precursor ions
           A <-
             ggplot(obj, aes(x = Ion.Inject.Time.in.ms)) +
             geom_histogram(binwidth = 1) +
             theme_bw() +
             xlab("Ion injection time [ms]")

           #precursor ion intensity
           B <-
             ggplot(obj, aes(x = Intensity)) +
             geom_histogram(bins = 100) +
             theme_bw() +
             scale_x_log10(labels = label_log(),
                           breaks = c(10^4, 10^5, 10^6, 10^7, 10^8, 10^9)) +
             xlab("Precursor intensity")

           #mass deviation
           C <-
             ggplot(obj, aes(x = Delta.M.in.ppm)) +
             geom_histogram(binwidth = 1) +
             geom_vline(xintercept = 0) +
             scale_x_continuous(breaks = seq(-10, 10, by = 2)) +
             theme_bw() +
             xlab("Mass deviation [ppm]")

           #mass deviation over retention time
           C1 <-
             ggplot(obj, aes(x = RT.in.min,
                             y = Delta.M.in.ppm)) +
             geom_point(fill = "grey25", alpha = 0.05) +
             xlim(0, 120) +
             geom_hline(yintercept = 0, colour = "red") +
             geom_hline(yintercept = c(-10, 10), colour = "red", linetype = "dashed") +
             theme_bw() +
             xlab("Retention time [min]")+
             ylab("Delta M [ppm]")

           #interference in isolation window
           D <-
             ggplot(obj, aes(x = Isolation.Interference.in.Percent)) +
```

```r
  geom_histogram(binwidth = 1) +
  geom_vline(xintercept = Isol_In,
             linetype = "dashed",
             colour = "red") +
  theme_bw() +
  xlab("Isolation interference [%]")

#percentage of SPS fragments matched to actual peptide fragments
E <-
  ggplot(obj, aes(x = SPS.Mass.Matches.in.Percent)) +
  geom_histogram(binwidth = 10) +
  geom_vline(xintercept = SPS_Mass_match - 5,
             linetype = "dashed",
             colour = "red") +
  scale_x_continuous(breaks = seq(0, 100, by = 10)) +
  theme_bw() +
  xlab("SPS Mass match [%]")

#signal-to-noise ratio of reporter ions
F1 <-
  ggplot(obj, aes(x = Average.Reporter.SN)) +
  geom_histogram(binwidth = 0.1) +
  geom_vline(xintercept = SN,
             linetype = "dashed",
             colour = "red") +
  theme_bw() +
  scale_x_log10(labels = label_log()) +
  annotation_logticks(sides = "b", base = 10) +
  xlab("Average reporter ion S/N ratio")

#average S/N threshold for PSM selection vs signal across channels
G <-
  obj %>%
  pivot_longer(cols = any_of(TMT_order),
               names_to = "Tag",
               values_to = "Abundance") %>%
  mutate(Tag = factor(x = Tag, levels = TMT_order)) %>%
  ggplot(aes(x = Tag, y = Abundance)) +
  geom_bin2d(bins = 60, drop = TRUE) +
  scale_fill_gradient(low = "grey25", high = "white") +
  theme_bw() +
  theme(axis.text.x = element_text(angle = 90, hjust = 1),
        legend.position = "bottom") +
  scale_y_log10(labels = trans_format(trans = "log10",
                                      format = math_format(10^.x)),
                breaks = c(0.1, 1, 10, 100, 1000, 10000)) +
  geom_hline(yintercept = SN,
             linetype = "dashed",
             colour = "red") +
  xlab("Reporter Ion") +
  ylab("Abundance [S/N]")

#plot all graphs as grid
```

```r
           plots <- plot_grid(A, B, C, C1, D, E, F1, G,
                              nrow = 2,
                              ncol = 4,
                              labels = "AUTO")

           #add title to each plot
           title <-
             ggdraw() +
             draw_label(label = paste(i,
                                      "PSMs N =",
                                      obj %>% nrow(),
                                      sep = " "),
                        fontface = 'bold',
                        x = 0,
                        hjust = 0) +
             theme(plot.margin = margin(0, 0, 0, 7))

           graphs_qual <-
             plot_grid(title,
                       plots,
                       ncol = 1,
                       rel_heights = c(0.1, 1))

           return(graphs_qual)

         }) %>%
  set_names(nm_batches)
```

```
## Warning: Removed 3 rows containing non-finite values (‘stat_bin()‘).

## Warning: Removed 235 rows containing non-finite values (‘stat_bin()‘).

## Warning: Removed 5600 rows containing non-finite values (‘stat_bin2d()‘).

## Warning: Removed 2 rows containing non-finite values (‘stat_bin()‘).

## Warning: Removed 174 rows containing non-finite values (‘stat_bin()‘).

## Warning: Removed 5304 rows containing non-finite values (‘stat_bin2d()‘).
```

```r
#save PSM quality graphs
#pdf(file = "graphs/PSM quality.pdf", width = 16, height = 8)
graphs_qual
```

```
## $Batch_1
```

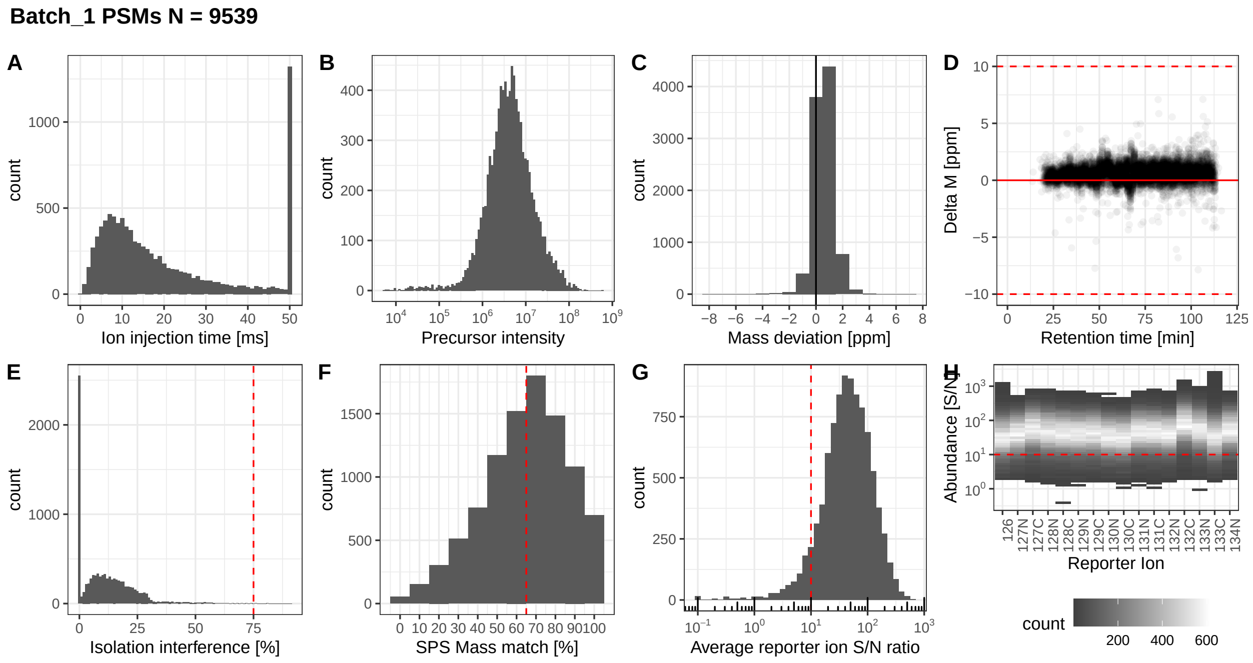


```
##
## $Batch_2
```

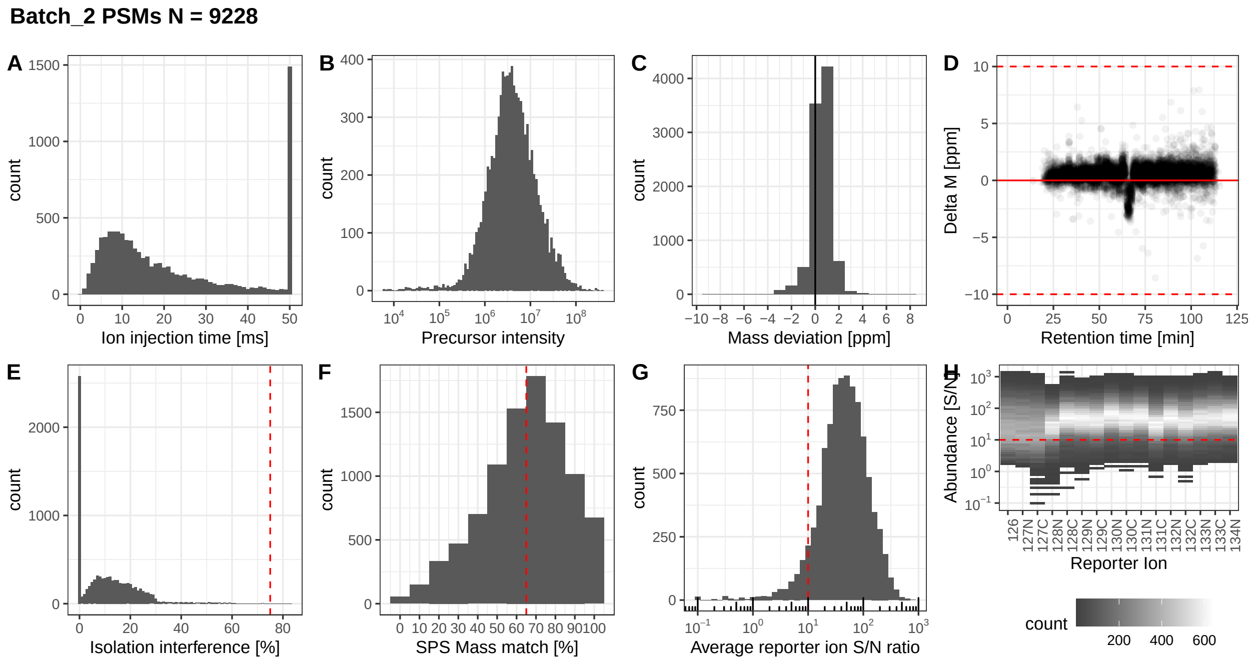


```
#dev.off()

#Sample loading (total intensity) between channels
#pdf(file = "graphs/raw total_signal.pdf", width = 16)
lapply(X = names(ls_Batch_PSM),
```

```r
FUN = function(i) {

  obj <- ls_Batch_PSM[[i]]

  Z <-
    obj %>%
    summarise(across(any_of(TMT_order),
                     ~ sum(.x, na.rm = TRUE))) %>%
    pivot_longer(cols = any_of(TMT_order),
                 names_to = "TMT.channel",
                 values_to = "total.intensity") %>%
    mutate(TMT.channel = factor(x = TMT.channel,
                                levels = TMT_order))

  plot <-
    ggplot(data = Z, aes(x = TMT.channel,
                         y = total.intensity)) +
    geom_col() +
    theme_bw() +
    xlab("TMT channel") +
    ylab("total reporter ion intensity") +
    labs(title = paste0(i))

  return(plot)
})
```

```
## [[1]]
```

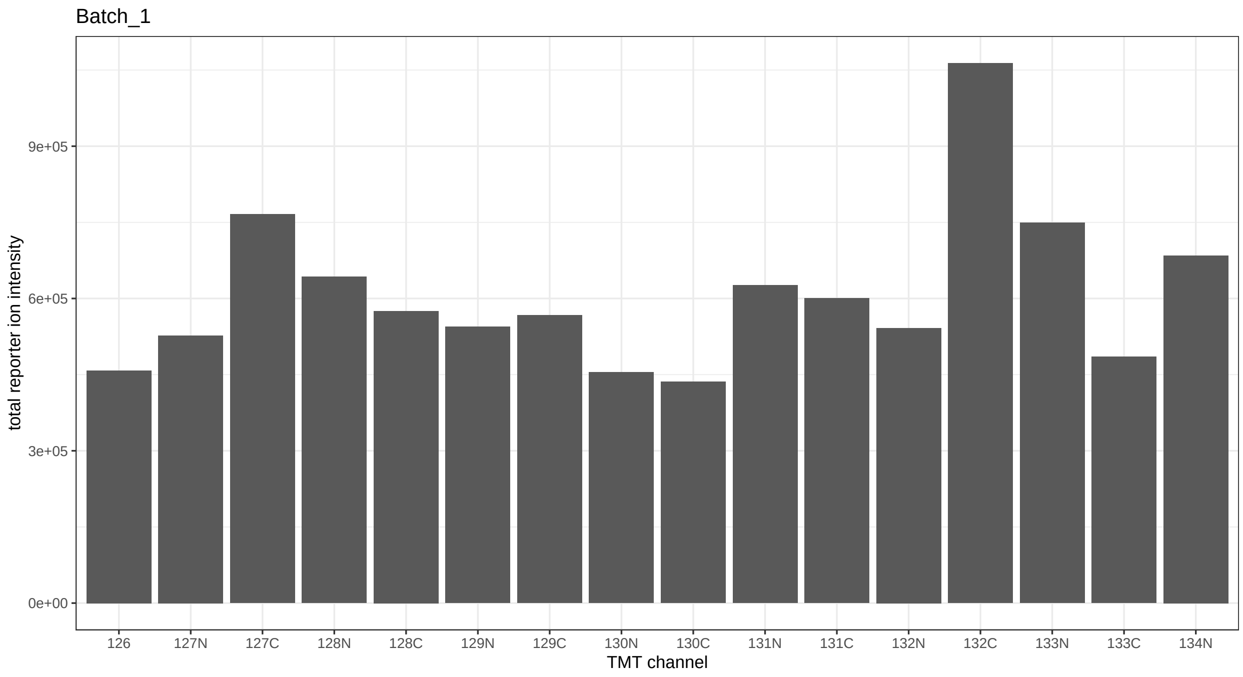


```
##
## [[2]]
```

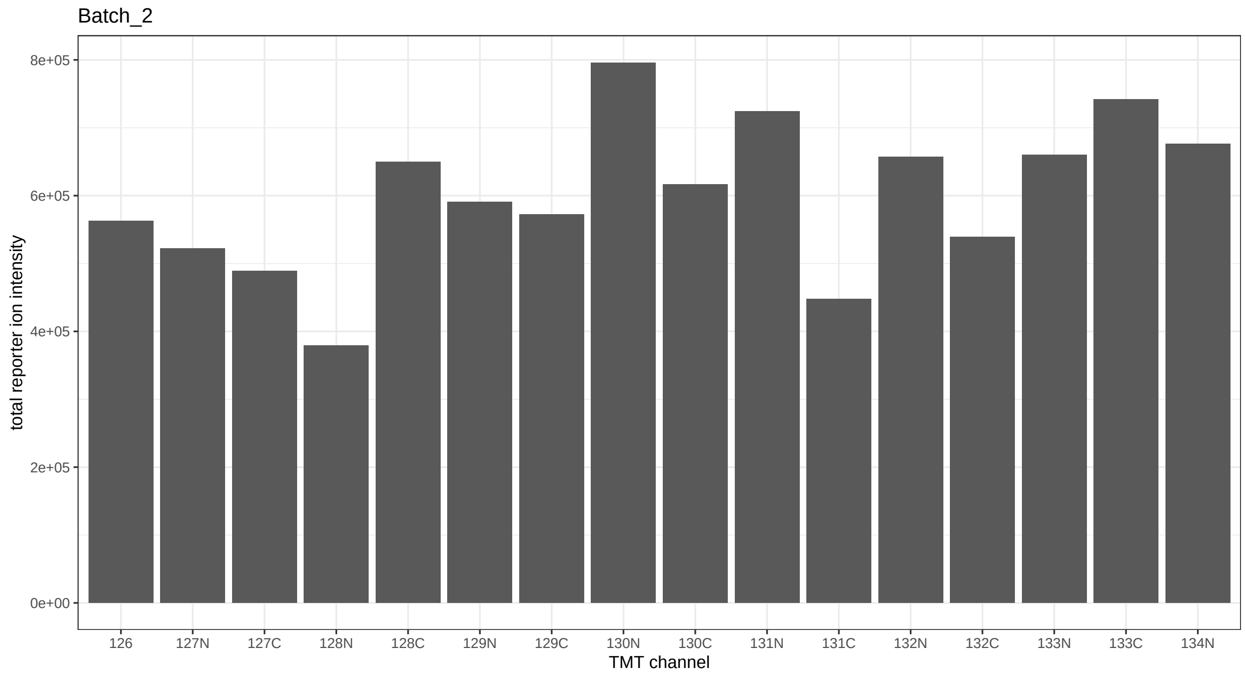


```r
#dev.off()


#overview of PSM assignments to Human and/or Toxoplasma database
ls_Batch_PSM[["Batch_1"]]$Marked.as %>% table()
```

```
## .
##             Human Human;Toxoplasma       Toxoplasma
##              2973              173             6297
```

```r
ls_Batch_PSM[["Batch_2"]]$Marked.as %>% table()
```

```
## .
##             Human Human;Toxoplasma       Toxoplasma
##              4237              146             4743
```

```r
#filtering of PSM data
ls_Batch_PSM_filt <-
  lapply(X = names(ls_Batch_PSM),
         FUN = function(i) {

           #get object
           obj <- ls_Batch_PSM[[i]]

           #quality filtering
           obj1 <-
             obj %>%
             filter(Master.Protein.Accessions != "") %>%
             filter(Contaminant == "FALSE") %>%
             filter(Marked.as == "Toxoplasma") %>%
```

```
             filter(PSM.Ambiguity == "Unambiguous"|PSM.Ambiguity == "Selected") %>%
             filter(Number.of.Protein.Groups == 1) %>%
             filter(is.na(Quan.Info) | Quan.Info != "NoQuanLabels") %>%
             filter(Concatenated.Rank == 1) %>%
             filter(Isolation.Interference.in.Percent <= Isol_In) %>%
             filter(SPS.Mass.Matches.in.Percent >= SPS_Mass_match) %>%
             filter(Average.Reporter.SN >= SN)

           return(obj1)
         }) %>%
  set_names(nm_batches)

rm(Isol_In, SPS_Mass_match, SN)
```

```
#creating Summarized Experiments for each cell line and adding metadata on replicate ID and condition (

####TgLMBD3
Annuli_TgLMBD3 <-
  readSummarizedExperiment(ls_Batch_PSM_filt[["Batch_1"]],
                           ecol = c("126", "127N", "127C",
                                    "128N", "128C", "129N"),
                           name = "psms") %>%
  zeroIsNA()

#sample layout
sample_layout_TgLMBD3 <-
  data.frame(TMT.channel = colnames(Annuli_TgLMBD3 %>% assay()),
             replicate = rep(c("rep1", "rep2", "rep3"), times = 2),
             condition = rep(c("KD", "control"), each = 3)) %>%
  mutate(sample.ID = paste(.$condition,
                           .$replicate,
                           sep = "_"))

sample_layout_TgLMBD3
```

```
##   TMT.channel replicate condition    sample.ID
## 1         126      rep1        KD      KD_rep1
## 2        127N      rep2        KD      KD_rep2
## 3        127C      rep3        KD      KD_rep3
## 4        128N      rep1   control control_rep1
## 5        128C      rep2   control control_rep2
## 6        129N      rep3   control control_rep3
```

```
#switching TMT channel names for sample IDs
colnames(Annuli_TgLMBD3) <- paste0(sample_layout_TgLMBD3$sample.ID)

#adding metadata to SumExp
Annuli_TgLMBD3$TMT.label <- sample_layout_TgLMBD3$TMT.channel
Annuli_TgLMBD3$condition <- sample_layout_TgLMBD3$condition
Annuli_TgLMBD3$replicate <- sample_layout_TgLMBD3$replicate
Annuli_TgLMBD3$sample.ID <- sample_layout_TgLMBD3$sample.ID
```

```r
#metadata in SumExp
colData(Annuli_TgLMBD3)
```

```
## DataFrame with 6 rows and 4 columns
##                TMT.label   condition   replicate     sample.ID
##              <character> <character> <character>   <character>
## KD_rep1              126          KD        rep1       KD_rep1
## KD_rep2             127N          KD        rep2       KD_rep2
## KD_rep3             127C          KD        rep3       KD_rep3
## control_rep1        128N     control        rep1 control_re...
## control_rep2        128C     control        rep2 control_re...
## control_rep3        129N     control        rep3 control_re...
```

```r
####TgNPSN
Annuli_TgNPSN <-
  readSummarizedExperiment(ls_Batch_PSM_filt[["Batch_1"]],
                           ecol = c("129C", "130N", "130C",
                                    "131N", "131C", "132N"),
                           name = "psms") %>%
  zeroIsNA()

Annuli_TgNPSN$SampleName <- c("TgNPSN_KD1","TgNPSN_KD2", "TgNPSN_KD3",
                              "TgNPSN_C1", "TgNPSN_C2", "TgNPSN_C3")
Annuli_TgNPSN$Condition <- rep(c("KD", "control"), each = 3)

#sample layout
sample_layout_TgNPSN <-
  data.frame(TMT.channel = colnames(Annuli_TgNPSN %>% assay()),
             replicate = rep(c("rep1", "rep2", "rep3"), times = 2),
             condition = rep(c("KD", "control"), each = 3)) %>%
  mutate(sample.ID = paste(.$condition,
                           .$replicate,
                           sep = "_"))

sample_layout_TgNPSN
```

```
##   TMT.channel replicate condition    sample.ID
## 1        129C      rep1        KD      KD_rep1
## 2        130N      rep2        KD      KD_rep2
## 3        130C      rep3        KD      KD_rep3
## 4        131N      rep1   control control_rep1
## 5        131C      rep2   control control_rep2
## 6        132N      rep3   control control_rep3
```

```r
#switching TMT channel names for sample IDs
colnames(Annuli_TgNPSN) <- paste0(sample_layout_TgNPSN$sample.ID)

#adding metadata to SumExp
Annuli_TgNPSN$TMT.label <- sample_layout_TgNPSN$TMT.channel
Annuli_TgNPSN$condition <- sample_layout_TgNPSN$condition
Annuli_TgNPSN$replicate <- sample_layout_TgNPSN$replicate
Annuli_TgNPSN$sample.ID <- sample_layout_TgNPSN$sample.ID
```

```r
#metadata in SumExp
colData(Annuli_TgNPSN)
```

```
## DataFrame with 6 rows and 6 columns
##                SampleName   Condition   TMT.label   condition   replicate
##               <character> <character> <character> <character> <character>
## KD_rep1       TgNPSN_KD1          KD        129C          KD        rep1
## KD_rep2       TgNPSN_KD2          KD        130N          KD        rep2
## KD_rep3       TgNPSN_KD3          KD        130C          KD        rep3
## control_rep1   TgNPSN_C1     control        131N     control        rep1
## control_rep2   TgNPSN_C2     control        131C     control        rep2
## control_rep3   TgNPSN_C3     control        132N     control        rep3
##                  sample.ID
##                <character>
## KD_rep1            KD_rep1
## KD_rep2            KD_rep2
## KD_rep3            KD_rep3
## control_rep1 control_re...
## control_rep2 control_re...
## control_rep3 control_re...
```

```r
####TgStxPM
Annuli_TgStxPM <-
  readSummarizedExperiment(ls_Batch_PSM_filt[["Batch_2"]],
                           ecol = c("126", "127N", "127C",
                                    "128N", "128C", "129N"),
                           name = "psms") %>%
  zeroIsNA()

#sample layout
sample_layout_TgStxPM <-
  data.frame(TMT.channel = colnames(Annuli_TgStxPM %>% assay()),
             replicate = rep(c("rep1", "rep2", "rep3"), times = 2),
             condition = rep(c("KD", "control"), each = 3)) %>%
  mutate(sample.ID = paste(.$condition,
                           .$replicate,
                           sep = "_"))

sample_layout_TgStxPM
```

```
##   TMT.channel replicate condition    sample.ID
## 1         126      rep1        KD      KD_rep1
## 2        127N      rep2        KD      KD_rep2
## 3        127C      rep3        KD      KD_rep3
## 4        128N      rep1   control control_rep1
## 5        128C      rep2   control control_rep2
## 6        129N      rep3   control control_rep3
```

```r
#switching TMT channel names for sample IDs
colnames(Annuli_TgStxPM) <- paste0(sample_layout_TgStxPM$sample.ID)

Annuli_TgStxPM$TMT.label <- sample_layout_TgStxPM$TMT.channel
```

```r
Annuli_TgStxPM$condition <- sample_layout_TgStxPM$condition
Annuli_TgStxPM$replicate <- sample_layout_TgStxPM$replicate
Annuli_TgStxPM$sample.ID <- sample_layout_TgStxPM$sample.ID

#metadata in SumExp
colData(Annuli_TgStxPM)
```

```
## DataFrame with 6 rows and 4 columns
##                TMT.label   condition   replicate     sample.ID
##              <character> <character> <character>   <character>
## KD_rep1              126          KD        rep1       KD_rep1
## KD_rep2             127N          KD        rep2       KD_rep2
## KD_rep3             127C          KD        rep3       KD_rep3
## control_rep1        128N     control        rep1 control_re...
## control_rep2        128C     control        rep2 control_re...
## control_rep3        129N     control        rep3 control_re...
```

```r
####TgSyp7
Annuli_TgSyp7 <-
  readSummarizedExperiment(ls_Batch_PSM_filt[["Batch_2"]],
                          ecol = c("129C", "130N", "130C",
                                   "132C", "133N", "133C"),
                          name = "psms") %>%
  zeroIsNA()

#sample layout
sample_layout_TgSyp7 <-
  data.frame(TMT.channel = colnames(Annuli_TgSyp7 %>% assay()),
             cell.line = "TgSyp7",
             replicate = rep(c("rep1", "rep2", "rep3"), times = 2),
             condition = rep(c("KD", "control"), each = 3)) %>%
  mutate(sample.ID = paste(.$condition,
                          .$replicate,
                          sep = "_"))

sample_layout_TgSyp7
```

```
##   TMT.channel cell.line replicate condition    sample.ID
## 1        129C    TgSyp7      rep1        KD      KD_rep1
## 2        130N    TgSyp7      rep2        KD      KD_rep2
## 3        130C    TgSyp7      rep3        KD      KD_rep3
## 4        132C    TgSyp7      rep1   control control_rep1
## 5        133N    TgSyp7      rep2   control control_rep2
## 6        133C    TgSyp7      rep3   control control_rep3
```

```r
#switching TMT channel names for sample IDs
colnames(Annuli_TgSyp7) <- paste0(sample_layout_TgSyp7$sample.ID)

Annuli_TgSyp7$TMT.label <- sample_layout_TgSyp7$TMT.channel
Annuli_TgSyp7$condition <- sample_layout_TgSyp7$condition
Annuli_TgSyp7$replicate <- sample_layout_TgSyp7$replicate
Annuli_TgSyp7$sample.ID <- sample_layout_TgSyp7$sample.ID
```

```r
#metadata in SumExp
colData(Annuli_TgSyp7)
```

```
## DataFrame with 6 rows and 4 columns
##                  TMT.label   condition   replicate     sample.ID
##                <character> <character> <character>   <character>
## KD_rep1              129C          KD        rep1       KD_rep1
## KD_rep2              130N          KD        rep2       KD_rep2
## KD_rep3              130C          KD        rep3       KD_rep3
## control_rep1         132C     control        rep1 control_re...
## control_rep2         133N     control        rep2 control_re...
## control_rep3         133C     control        rep3 control_re...
```

```r
#create a list of the Summarized Experiments
ls_SumExps <-
  list(Annuli_TgLMBD3,
       Annuli_TgNPSN,
       Annuli_TgStxPM,
       Annuli_TgSyp7) %>%
  set_names(nm_cell_lines)

rm(sample_layout_TgStxPM, sample_layout_TgNPSN, sample_layout_TgSyp7, sample_layout_TgLMBD3)
```

```r
#Overview of how many PSMs have how many missing TMT channels
#pdf(file = "graphs/PSM_MV_pattern.pdf", width = 11, height = 8)
lapply(X = names(ls_SumExps),
       FUN = function(i) {

         #get object
         obj <- ls_SumExps[[i]]

         #Pattern of missing PSMs per channel
         A <-
           #create dataframe with channel name and percentage of missing values
           data.frame(nNA(obj) %>% .$nNAcols %>% .$name,
                      nNA(obj) %>% .$nNAcols %>% .$pNA) %>%
           setNames(c("channel", "pNA")) %>%
           #plot
           ggplot(aes(x = factor(channel,
                                 levels = channel),
                      y = pNA)) +
           geom_col() +
           geom_text(aes(label = round(pNA, 3)),
                     vjust = -0.2) +
           ggtitle("Proportion of PSMs with missing quantitation", i) +
           xlab("sample ID") +
           ylab("% missing") +
           theme_bw() +
           scale_x_discrete(guide = guide_axis(angle = 45))

         #How many PSMs have how many missing TMT channels
         B <-
```

```
        nNA(obj) %>% .$nNArows %>% .$nNA %>% table() %>% as.data.frame() %>%
        mutate(share = round(Freq/nrow(obj) *100, digits = 1)) %>%
        set_colnames(c("No.of.missing.channels", "Count", "Freq")) %>%
        #plot
        ggplot(aes(x = No.of.missing.channels,
                   y = Count)) +
        geom_col() +
        geom_text(aes(label = paste0(Freq, "%")),
                  vjust = -0.2) +
        ggtitle("PSMs with missing channels [%]",
                paste(i, "total:", obj %>% nrow(), sep = " ")) +
        xlab("total number of missing channels") +
        ylab("N") +
        ylim(0, nrow(obj)) +
        theme_bw()

      Z <-
        plot_grid(A, B,
                  nrow = 1,
                  ncol = 2,
                  labels = "AUTO")
      return(Z)
    })
```

```
## [[1]]
```

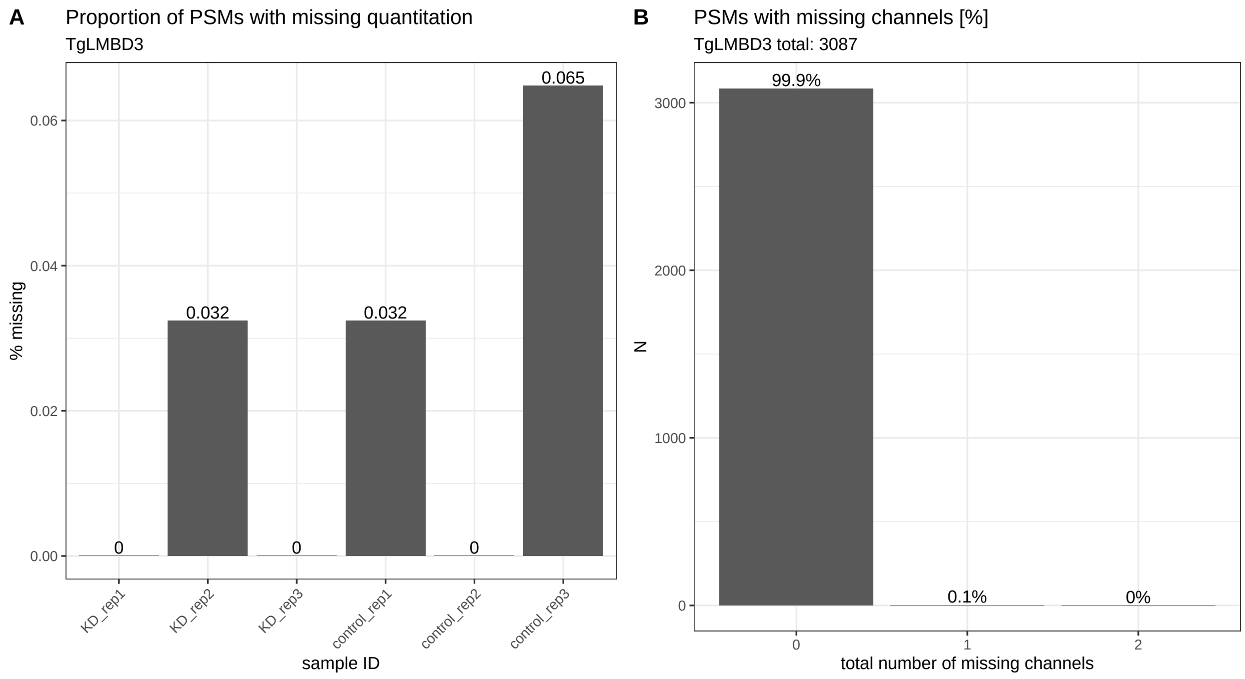


```
##
## [[2]]
```

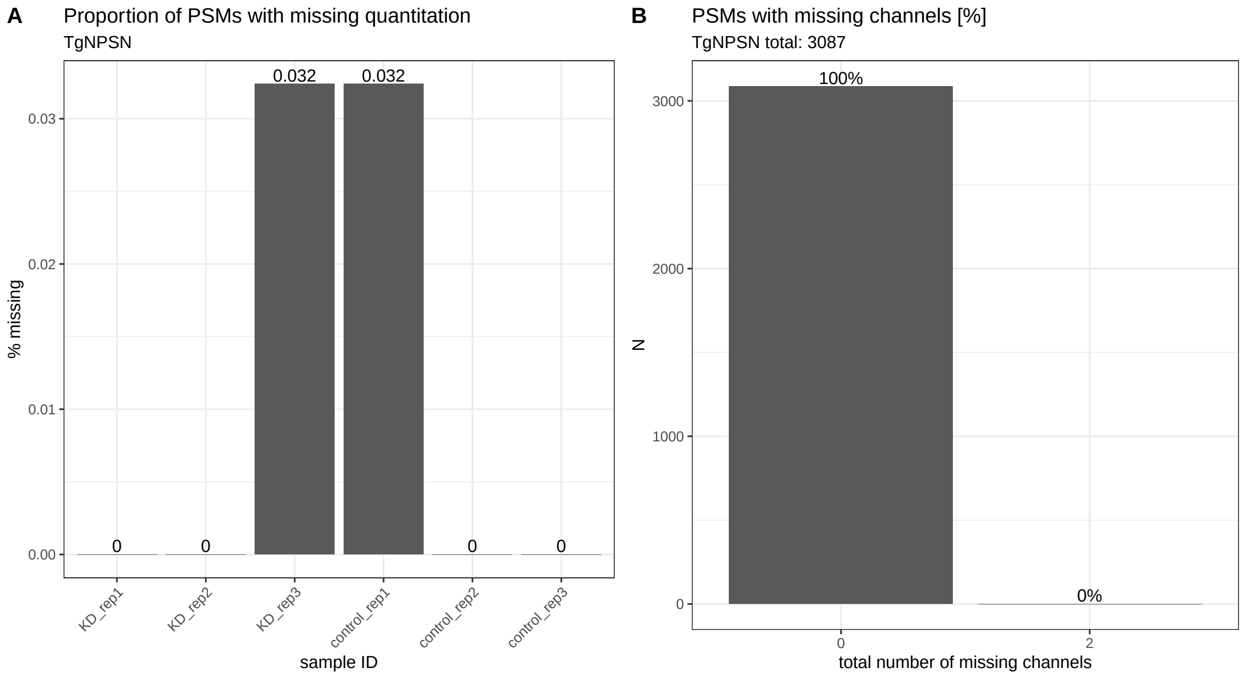
A
Proportion of PSMs with missing quantitation
TgNPSN
0.032
0.032
0
0
0
0
0.03
0.02
0.01
0.00
% missing
KD_rep1
KD_rep2
KD_rep3
control_rep1
control_rep2
control_rep3
sample ID
B
PSMs with missing channels [%]
TgNPSN total: 3087
100%
0%
3000
2000
1000
0
N
0
2
total number of missing channels

```
##
## [[3]]
```

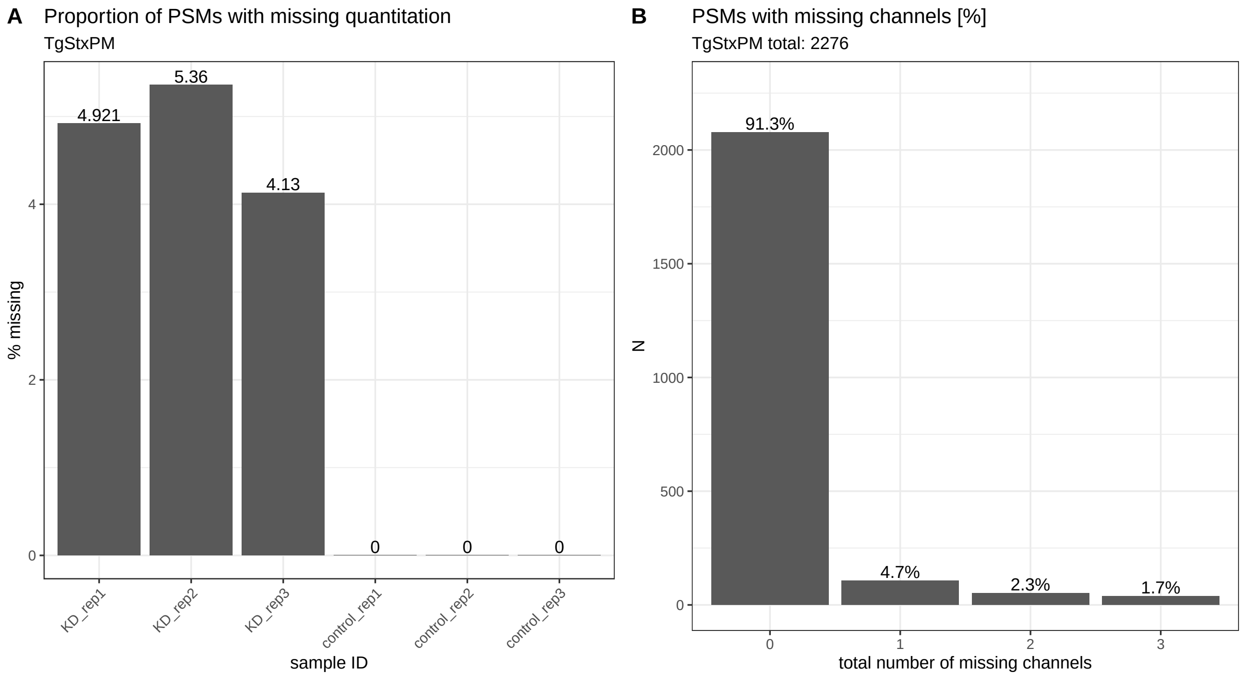
A
Proportion of PSMs with missing quantitation
TgStxPM
5.36
4.921
4.13
0
0
0
4
2
0
% missing
KD_rep1
KD_rep2
KD_rep3
control_rep1
control_rep2
control_rep3
sample ID
B
PSMs with missing channels [%]
TgStxPM total: 2276
91.3%
4.7%
2.3%
1.7%
2000
1500
1000
500
0
N
0
1
2
3
total number of missing channels

```
##
## [[4]]
```

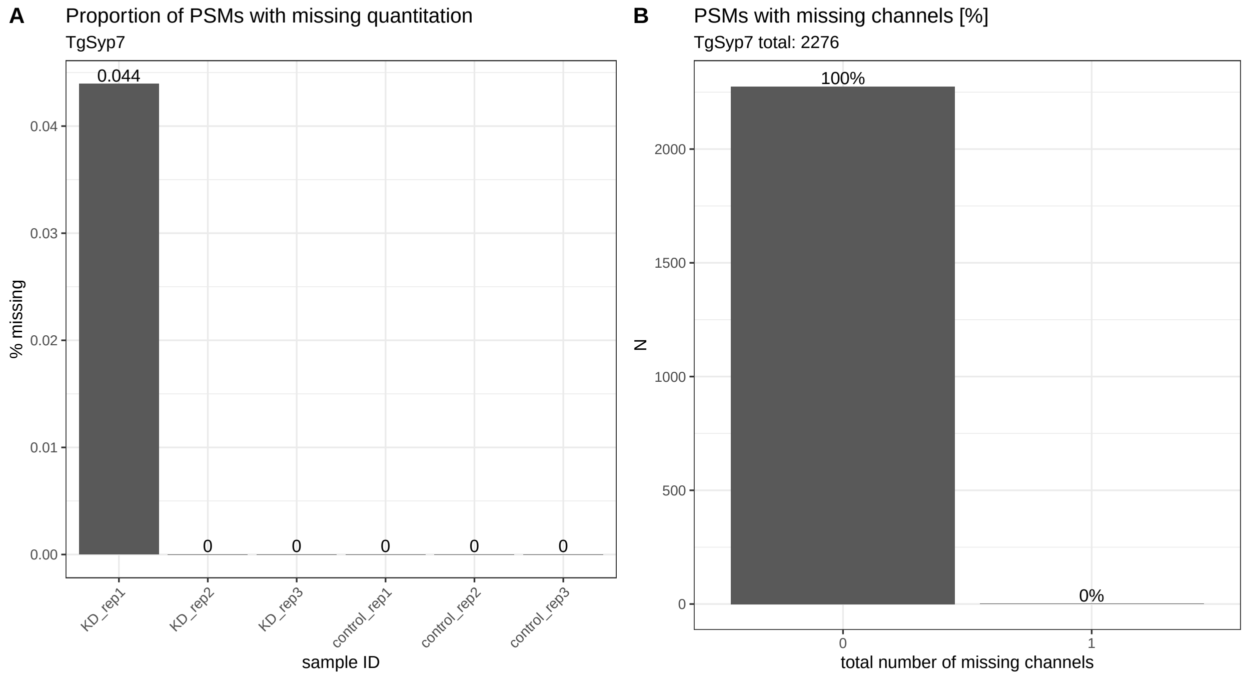


```r
#dev.off()

#Reduction of PSM datasets to consider only PSMs with no missing values
ls_SumExps_complete <-
  lapply(X = names(ls_SumExps),
         FUN = function(i) {

           #get object
           obj <- ls_SumExps[[i]]

           #remove PSMs that have missing values
           obj <- filterNA(obj, pNA = 0)

           return(obj)
         }) %>%
  set_names(nm_cell_lines)
```

```r
#Aggregation from psm to protein level
ls_QFeat <-
  lapply(X = names(ls_SumExps_complete),
         FUN = function(i) {

           #get object
           obj <- ls_SumExps_complete[[i]]

           #read Summarized Experiment into a QFeatures object
           QFeat <- QFeatures(List(psms = obj))

           #re-add colData
           colData(QFeat) <- colData(obj)
```

```
          ##log2 transform psm data for robustSummary aggregation
          QFeat1 <- logTransform(QFeat,
                                 base = 2,
                                 i = "psms",
                                 name = "log2_psms")

          ##aggregation psm to peptide using iterated re-weighted least squares (IWLS)
          QFeat2 <- aggregateFeatures(object = QFeat1,
                                      i = "log2_psms",
                                      fcol = "Sequence",
                                      name = "log2_peptides",
                                      fun = MsCoreUtils::robustSummary)

          #aggregation from psm to protein using iterated re-weighted least squares (IWLS)
          QFeat3 <- aggregateFeatures(object = QFeat2,
                                      i = "log2_peptides",
                                      fcol = "Master.Protein.Accessions",
                                      name = "log2_proteins",
                                      fun = MsCoreUtils::robustSummary)

          return(QFeat3)
        }) %>%
  set_names(nm_cell_lines)
```

```
####removal of low FDR proteins using PD information
ls_QFeat <-
  lapply(X = names(ls_QFeat),
         FUN = function(i) {

          obj <- ls_QFeat[[i]]

          #protein file
          df_prot <-
            read_delim(file = paste0("input/P1218_Toxo_Annuli_Proteins.txt"),
                       delim = "\t",
                       na = c("", "NA"),
                       col_types = NULL)

          #vector of QFeat accessions with high (FDR 1%) and medium (FDR 5%) protein confidence
          vct_confid_accessions <-
            df_prot %>%
            filter(`Protein FDR Confidence Combined` %in% c("High", "Medium")) %>%
            filter(Accession %in% rownames(ls_QFeat[[i]][["log2_proteins"]])) %>%
            .$Accession

          #creating SumExp object containing confident accession numbers
          QFeat_confid <- ls_QFeat[[i]][["log2_proteins"]][vct_confid_accessions]

          #add SumExp with confident proteins to QFeature object
          obj1 <- addAssay(obj,
                           QFeat_confid,
                           name = "log2_proteins_FDR5")
```

```r
          #add extra assay to fill with untransformed protein values for normalyzer test
          obj2 <- addAssay(obj1,
                           obj1[["log2_proteins_FDR5"]],
                           name = "proteins_FDR5")

          #replace assay data in "proteins" layer with anti-log2 values
          assay(obj2[["proteins_FDR5"]]) <- 2^(assay(obj2[["log2_proteins_FDR5"]]))+1 #+1 to avoid iss

          message(paste0("QFeatures structure: ", i))

          print(obj2)

          return(obj2)
        }) %>%
  set_names(nm_cell_lines)
```

```
## An instance of class QFeatures containing 6 assays:
##  [1] psms: SummarizedExperiment with 3084 rows and 6 columns
##  [2] log2_psms: SummarizedExperiment with 3084 rows and 6 columns
##  [3] log2_peptides: SummarizedExperiment with 2902 rows and 6 columns
##  [4] log2_proteins: SummarizedExperiment with 817 rows and 6 columns
##  [5] log2_proteins_FDR5: SummarizedExperiment with 817 rows and 6 columns
##  [6] proteins_FDR5: SummarizedExperiment with 817 rows and 6 columns
## An instance of class QFeatures containing 6 assays:
##  [1] psms: SummarizedExperiment with 3086 rows and 6 columns
##  [2] log2_psms: SummarizedExperiment with 3086 rows and 6 columns
##  [3] log2_peptides: SummarizedExperiment with 2903 rows and 6 columns
##  [4] log2_proteins: SummarizedExperiment with 817 rows and 6 columns
##  [5] log2_proteins_FDR5: SummarizedExperiment with 817 rows and 6 columns
##  [6] proteins_FDR5: SummarizedExperiment with 817 rows and 6 columns
## An instance of class QFeatures containing 6 assays:
##  [1] psms: SummarizedExperiment with 2078 rows and 6 columns
##  [2] log2_psms: SummarizedExperiment with 2078 rows and 6 columns
##  [3] log2_peptides: SummarizedExperiment with 1985 rows and 6 columns
##  [4] log2_proteins: SummarizedExperiment with 650 rows and 6 columns
##  [5] log2_proteins_FDR5: SummarizedExperiment with 650 rows and 6 columns
##  [6] proteins_FDR5: SummarizedExperiment with 650 rows and 6 columns
## An instance of class QFeatures containing 6 assays:
##  [1] psms: SummarizedExperiment with 2275 rows and 6 columns
##  [2] log2_psms: SummarizedExperiment with 2275 rows and 6 columns
##  [3] log2_peptides: SummarizedExperiment with 2179 rows and 6 columns
##  [4] log2_proteins: SummarizedExperiment with 672 rows and 6 columns
##  [5] log2_proteins_FDR5: SummarizedExperiment with 672 rows and 6 columns
##  [6] proteins_FDR5: SummarizedExperiment with 672 rows and 6 columns
```

```r
#normalyzer test on each dataset to see effect of different normalisation methods
lapply(X = names(ls_QFeat),
       function(i) {

         obj <- ls_QFeat[[i]]

         normalyzer(jobName = paste("normalyzer", i, sep = "_"),
                    experimentObj = obj[["proteins_FDR5"]],
```

```
                     sampleColName = "sample.ID",
                     groupColName = "condition",
                     outputDir = "files")
       })
```

```
## [[1]]
## NULL
##
## [[2]]
## NULL
##
## [[3]]
## NULL
##
## [[4]]
## NULL
```

```
#apply chosen normalization method across all samples
norm_method <- "center.median"

ls_QFeat <-
  lapply(X = names(ls_QFeat),
         FUN = function(i) {

           #get object
           obj <- ls_QFeat[[i]]

           #alignment of global protein abundances across all samples
           QFeat <- normalize(obj,
                               i = "log2_proteins_FDR5",
                               name = "log2_norm_proteins_FDR5",
                               method = norm_method) #center.median normalisation
           return(QFeat)
         }) %>%
  set_names(nm_cell_lines)


#save list of QFeat containers as importable file
saveRDS(object = ls_QFeat,
        file = "files/list_of_Toxo_annuli_QFeatures_datasets.rds")


rm(norm_method)

#Plots of intensity signal across data levels and samples
#pdf(file = "graphs/intensity distribution.pdf", width = 16, height = 8)
invisible(lapply(X = names(ls_QFeat),
       function(i){

         #get object
         obj <- ls_QFeat[[i]]

         #plot
```

```
    par(mfrow = c(2, 3), las = 2)
    boxplot(assay(obj[["psms"]]),
            main = paste(i, "raw psms", sep = " "),
            ylab = "intensity")
    boxplot(assay(obj[["log2_proteins"]]),
            main = paste(i, "log2 proteins", sep = " "),
            ylab = "log2 intensity")
    boxplot(assay(obj[["log2_norm_proteins_FDR5"]]),
            main = paste(i, "log2 proteins normalised; FDR 5%", sep = " "),
            ylab = "log2 intensity")
    plotDensities(assay(obj[["psms"]]),
                  legend = "topright",
                  group = obj$condition)
    plotDensities(assay(obj[["log2_proteins"]]),
                  legend = FALSE,
                  group = obj$condition)
    plotDensities(object = assay(obj[["log2_norm_proteins_FDR5"]]),
                  legend = FALSE,
                  group = obj$condition)
  }))
```

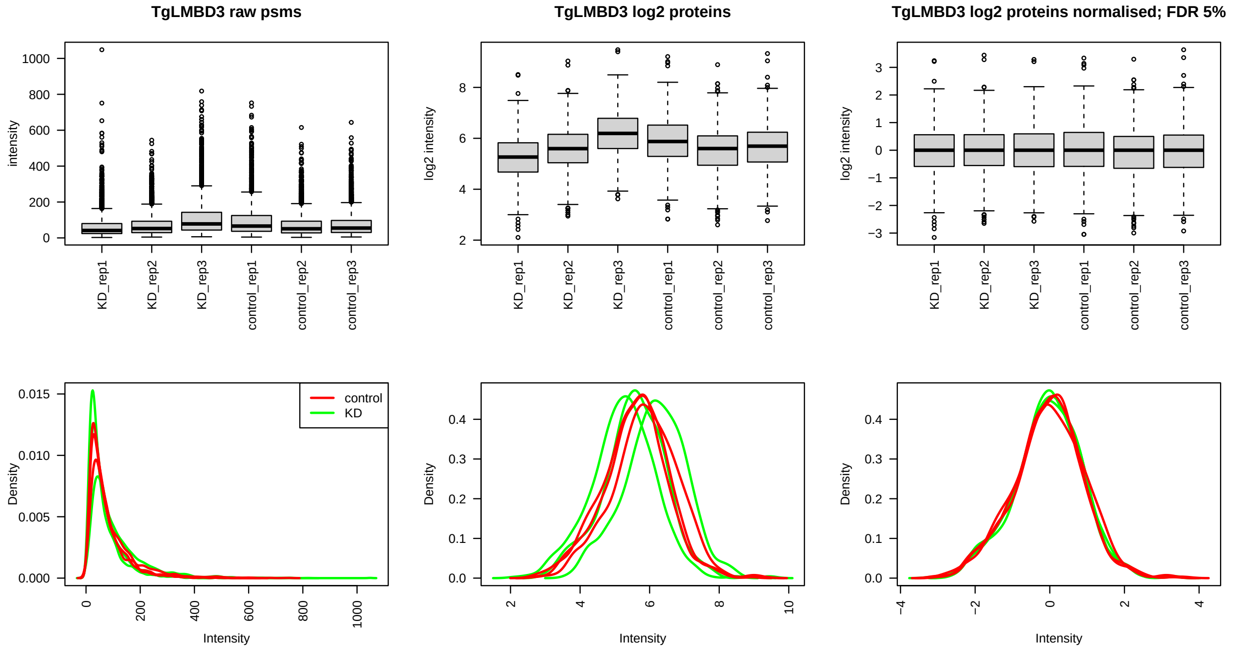

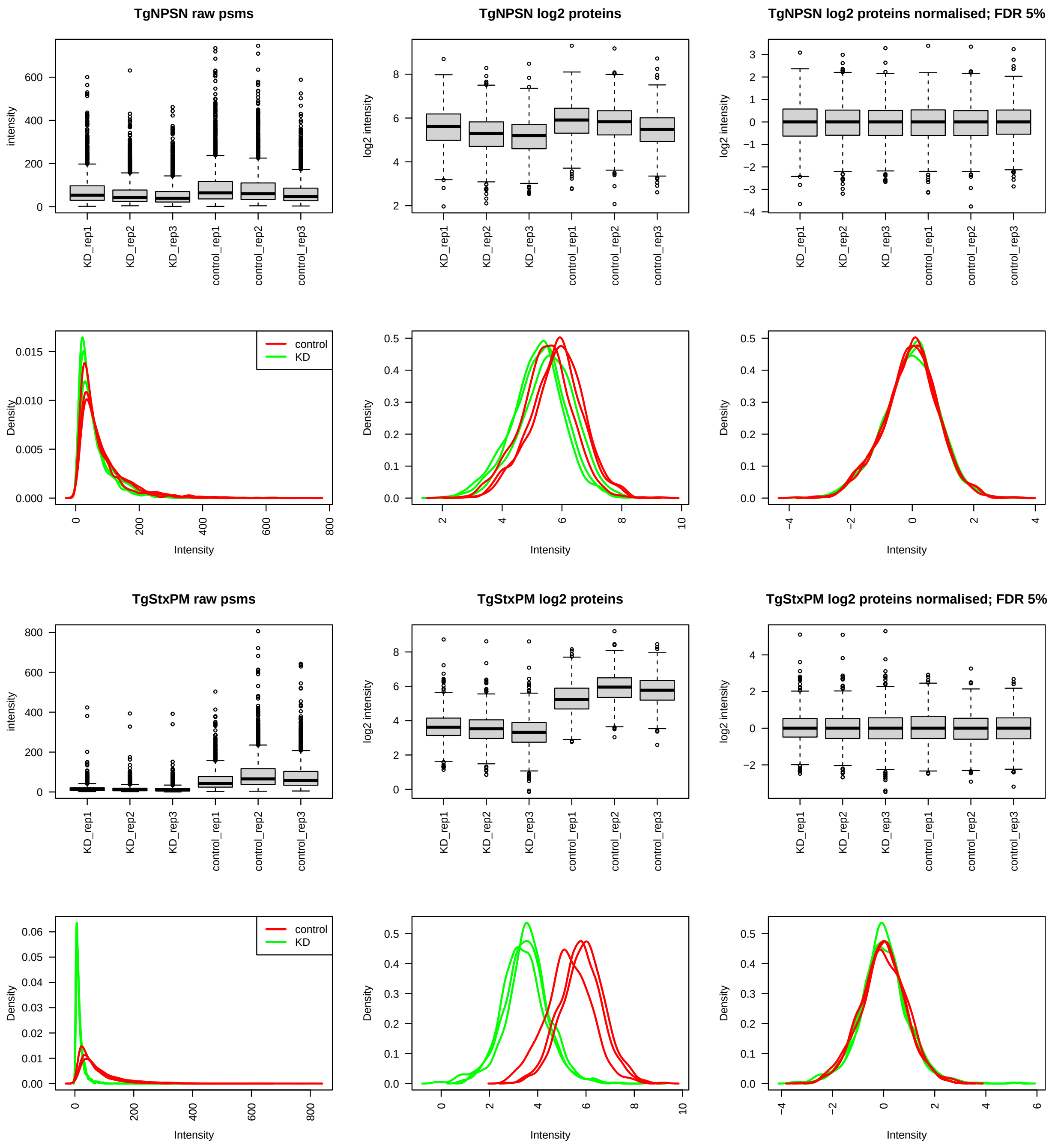
TgNPSN raw psms
TgNPSN log2 proteins
TgNPSN log2 proteins normalised; FDR 5%
intensity
log2 intensity
KD_rep1
KD_rep2
KD_rep3
control_rep1
control_rep2
control_rep3
control
KD
Density
Intensity
TgStxPM raw psms
TgStxPM log2 proteins
TgStxPM log2 proteins normalised; FDR 5%

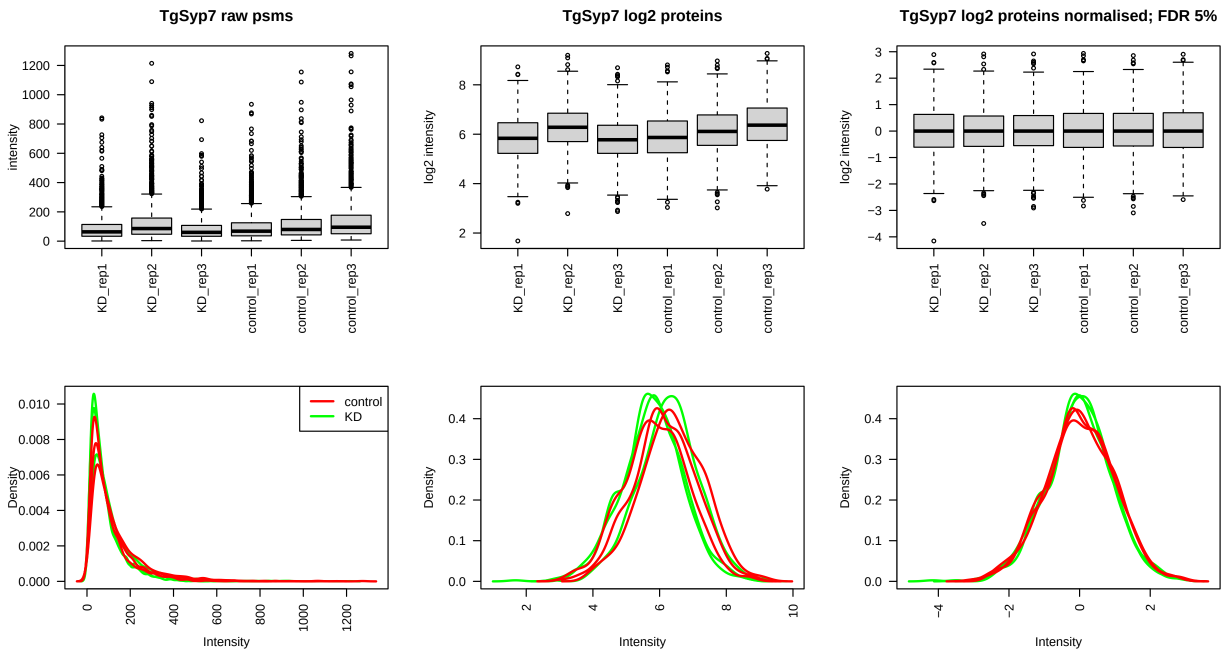


```
#dev.off()

#save untransformed and log2-transformed, normalized protein intensities to output file
invisible(lapply(X = names(ls_QFeat),
                 function(i) {

                   #get raw and transformed log2-normalized protein data
                   obj_raw <- ls_QFeat[[i]][["proteins_FDR5"]]
                   obj_final <- ls_QFeat[[i]][["log2_norm_proteins_FDR5"]]

                   #raw data - combining Master.Protein.Accessions, number of peptides, and assay data
                   obj_raw <-
                     cbind(rowData(ls_QFeat[[i]][["proteins_FDR5"]]) %>%
                             as.data.frame() %>%
                             select(Master.Protein.Accessions, .n) %>%
                             set_colnames(c("Accession", "number.of.peptides")),
                           assay(obj_raw))

                   write_csv(x = obj_raw,
                             file = paste0("files/", i, "_raw_protein_abundances.csv"))

                   obj_final <-
                     cbind(rowData(ls_QFeat[[i]][["log2_norm_proteins_FDR5"]]) %>%
                             as.data.frame() %>%
                             select(Master.Protein.Accessions, .n) %>%
                             set_colnames(c("Accession", "number.of.peptides")),
                           assay(obj_final))

                   write_csv(x = obj_final,
                             file = paste0("files/", i,
                                           "_FDR5_log2_median_normalised_protein_abundances.csv"))
```

```
                 }))

#list of master protein accession numbers in each set
Euler_list <-
  list(TgLMBD3 =
         rowData(ls_QFeat[["TgLMBD3"]][["log2_norm_proteins_FDR5"]])$Master.Protein.Accessions %>%
         unique(),
       TgNPSN =
         rowData(ls_QFeat[["TgNPSN"]][["log2_norm_proteins_FDR5"]])$Master.Protein.Accessions %>%
         unique(),
       TgStxPM =
         rowData(ls_QFeat[["TgStxPM"]][["log2_norm_proteins_FDR5"]])$Master.Protein.Accessions %>%
         unique(),
       TgSyp7 =
         rowData(ls_QFeat[["TgSyp7"]][["log2_norm_proteins_FDR5"]])$Master.Protein.Accessions %>%
         unique())

#Euler plot
#pdf(file = "graphs/Euler_plot_protein_overlap.pdf")
plot(euler(Euler_list),
     fill = c("#E69F00", "#009E73", "dodgerblue4"),
     quantities = TRUE)
```

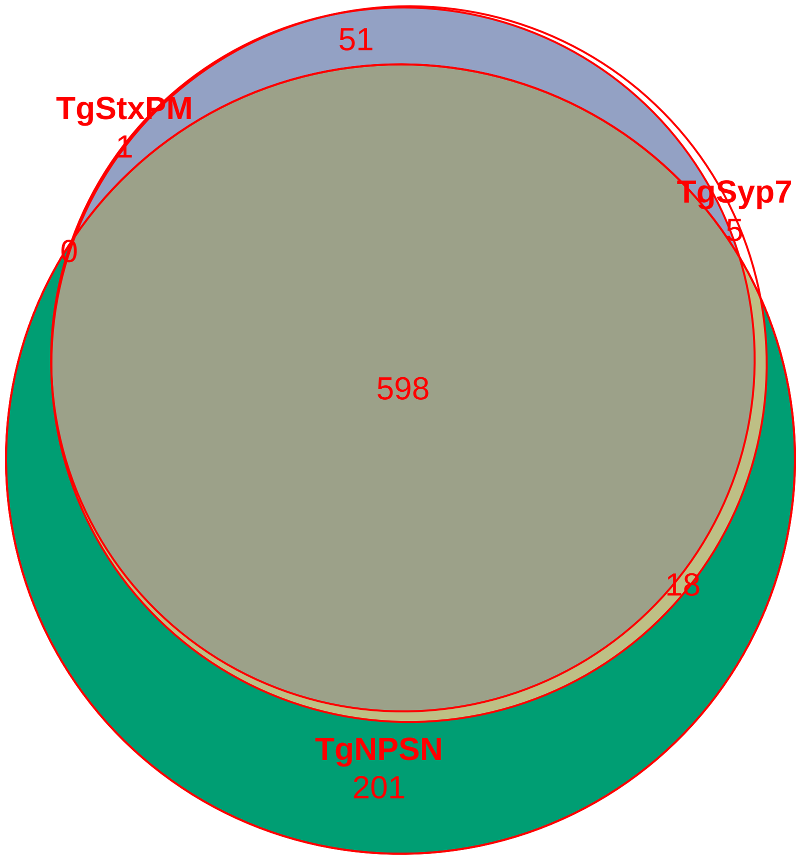

```r
#dev.off()

rm(Euler_list)
```

```r
#Protein PSM support pattern
#pdf(file = "graphs/protein_PSM_support.pdf", width = 11, height = 8)
lapply(X = names(ls_QFeat),
       FUN = function(i) {

         #get object
         obj <- ls_QFeat[[i]][["psms"]]

         #PSM count table for each master protein accession
         PSM_count <-
           obj %>% rowData() %>% .$Master.Protein.Accessions %>%
           table() %>%
           as.data.frame.table

         #summary table listing PSM per protein
         freq_table <-
           PSM_count %>%
           group_by(Freq) %>%
           tally()

         percent_table <-
           freq_table %>%
           #add new column, defining the grouping categories
           mutate(thresh = case_when(Freq <= 5 ~ as.character(Freq),
                                     Freq > 5 ~ "6+")) %>%
           group_by(thresh) %>%
           summarise(N = sum(n),
                     groups = levels(thresh)) %>%
           mutate(percentage = round((N/sum(N)) *100, digits = 1))

         #plot PSM support per protein
         bar_chart <-
           percent_table %>%
           ggplot(aes(x = thresh,
                      y = N)) +
           geom_col() +
           geom_text(aes(label = N),
                     vjust = -0.2) +
           ggtitle(paste("PSM support per protein:",
                         i,
                         "N =",
                         PSM_count$. %>% unique() %>% length(),
                         sep = " ")) +
           xlab("number of PSMs per protein") +
           ylab("N") +
           theme_bw()

         #plot relative proportion of PSM support categories
         pie_chart <-
```

```
      percent_table %>%
      ggplot(aes(x = "",
                 y = percentage,
                 fill = thresh)) +
      geom_col() +
      scale_fill_brewer(palette = "Greys",
                        direction = -1) +
      guides(fill = guide_legend(title = "# of PSMs")) +
      theme_void() +
      geom_text(aes(label = paste0(percentage, "%")),
                colour = "red",
                position = position_stack((vjust = 0.5))) +
      coord_polar(theta = "y",
                  start = 0,
                  direction = -1)

    plot <- plot_grid(bar_chart, pie_chart,
                      nrow = 1,
                      ncol = 2,
                      labels = "AUTO")
    return(plot)
  })
```

```
## [[1]]
```

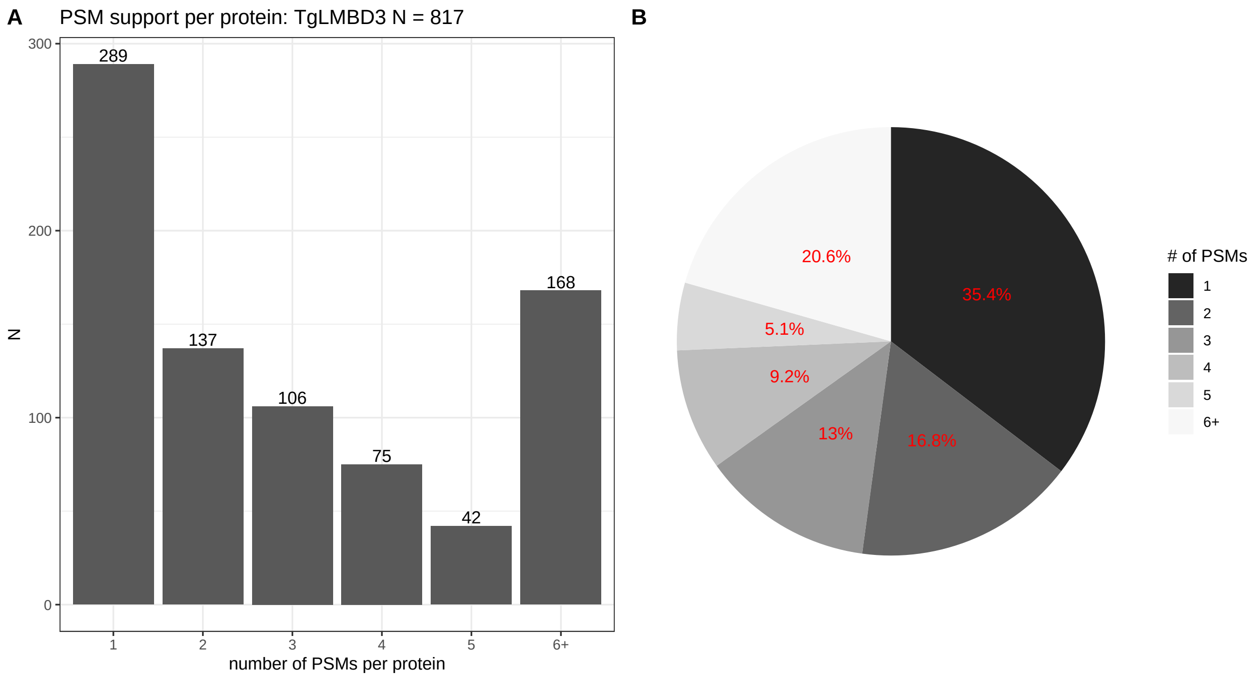


```
##
## [[2]]
```

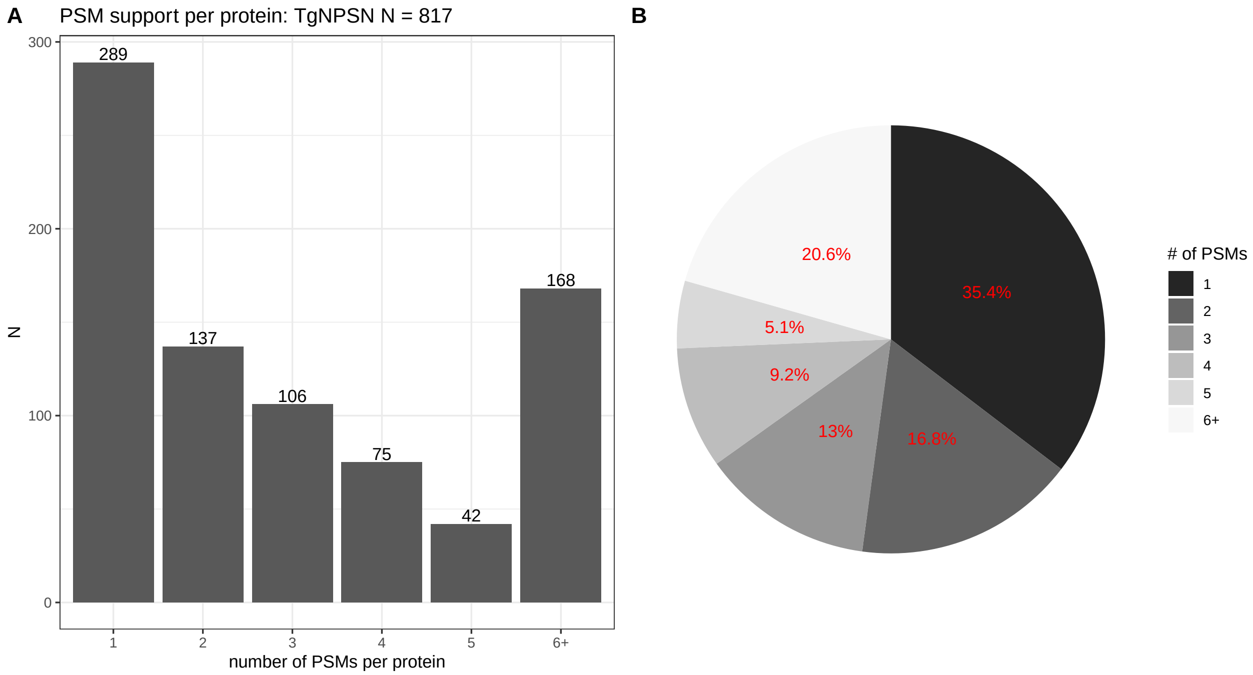
A
PSM support per protein: TgNPSN N = 817
B
N
number of PSMs per protein
289
137
106
75
42
168
0
100
200
300
1
2
3
4
5
6+
20.6%
35.4%
5.1%
9.2%
13%
16.8%
### of PSMs
1
2
3
4
5
6+

```
##
## [[3]]
```

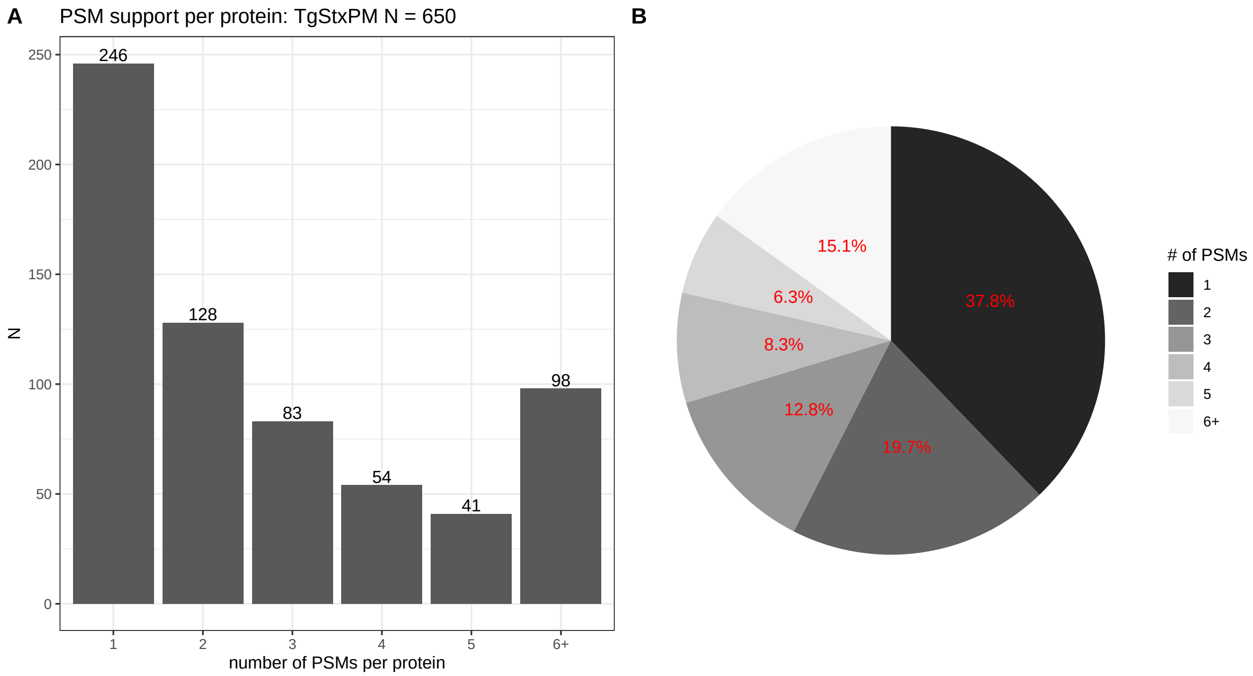
A
PSM support per protein: TgStxPM N = 650
B
N
number of PSMs per protein
246
128
83
54
41
98
0
50
100
150
200
250
1
2
3
4
5
6+
15.1%
37.8%
6.3%
8.3%
12.8%
19.7%
### of PSMs
1
2
3
4
5
6+

```
##
## [[4]]
```

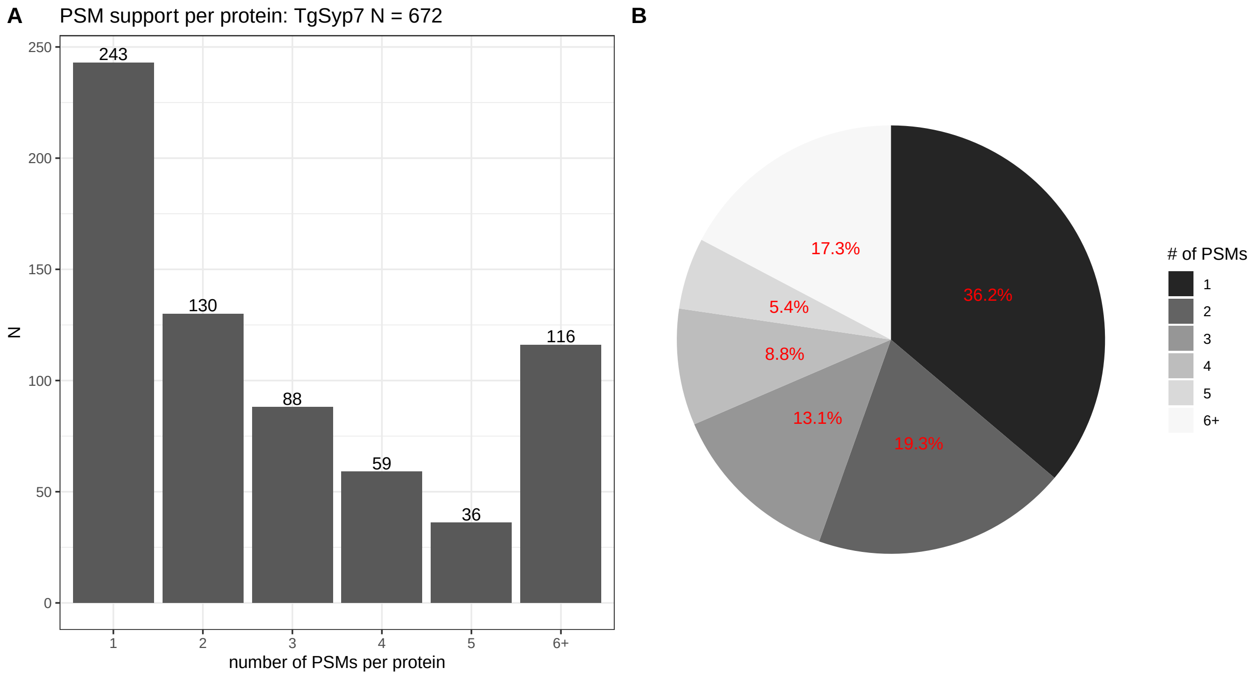


```r
#dev.off()
```

```r
ls_PCA <-
  lapply(X = names(ls_QFeat),
         FUN = function(i) {

           #get object
           obj <- ls_QFeat[[i]][["log2_norm_proteins_FDR5"]]

           PCA_obj <- PCA(X = assay(obj) %>% t(),
                          ncp = 10,
                          scale.unit = TRUE,
                          graph = FALSE)

           #Scree plot
           PCA_scree <-
             fviz_screeplot(PCA_obj,
                            choice = "variance",
                            addlabels = TRUE,
                            ncp = 8,
                            main = paste(i, "Scree plot", sep = " "))

           PCA_score_1_2 <-
             fviz_pca_ind(PCA_obj,
                          axes = c(1, 2),
                          geom = c("point", "text"),
                          habillage = colData(obj)$condition %>% as.factor(),
                          title = paste(i, "PC1_PC2", sep = " "),
                          addEllipses = FALSE,
                          repel = TRUE) +
             theme_bw()
```

```r
          PCA_score_1_3 <-
            fviz_pca_ind(PCA_obj,
                         axes = c(1, 3),
                         geom = c("point", "text"),
                         habillage = colData(obj)$condition %>% as.factor(),
                         title = paste(i, "PC1_PC3", sep = " "),
                         addEllipses = FALSE,
                         repel = TRUE) +
            theme_bw()

          PCA_score_2_3 <-
            fviz_pca_ind(PCA_obj,
                         axes = c(2, 3),
                         geom = c("point", "text"),
                         habillage = colData(obj)$condition %>% as.factor(),
                         title = paste(i, "PC2_PC3", sep = " "),
                         addEllipses = FALSE,
                         repel = TRUE) +
            theme_bw()

          plot <-
            plot_grid(PCA_scree, PCA_score_1_2, PCA_score_1_3, PCA_score_2_3,
                      nrow = 2, ncol = 2,
                      labels = "AUTO",
                      greedy = TRUE)

          items <- list(PCA_object = PCA_obj,
                        PCA_plots = plot)

          return(items)
        }) %>%
  set_names(nm_cell_lines)

#save all plots
#pdf(file = "graphs/PCA_plots.pdf", width = 11, height = 8)
for(i in seq(1, length(ls_PCA))) {
  print(ls_PCA[[i]][["PCA_plots"]])
}
```

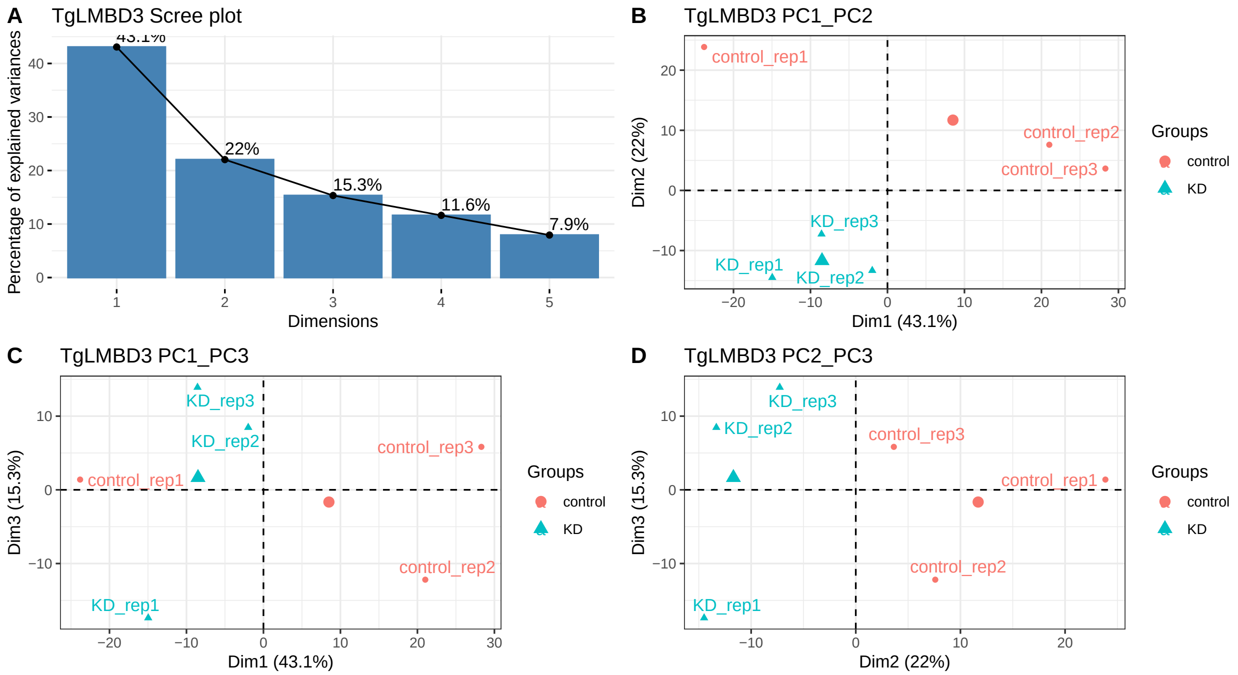

A
TgLMBD3 Scree plot
Percentage of explained variances
Dimensions
43.1%
22%
15.3%
11.6%
7.9%
B
TgLMBD3 PC1_PC2
Dim1 (43.1%)
Dim2 (22%)
control_rep1
control_rep2
control_rep3
KD_rep1
KD_rep2
KD_rep3
Groups
control
KD
C
TgLMBD3 PC1_PC3
Dim1 (43.1%)
Dim3 (15.3%)
D
TgLMBD3 PC2_PC3
Dim2 (22%)
Dim3 (15.3%)


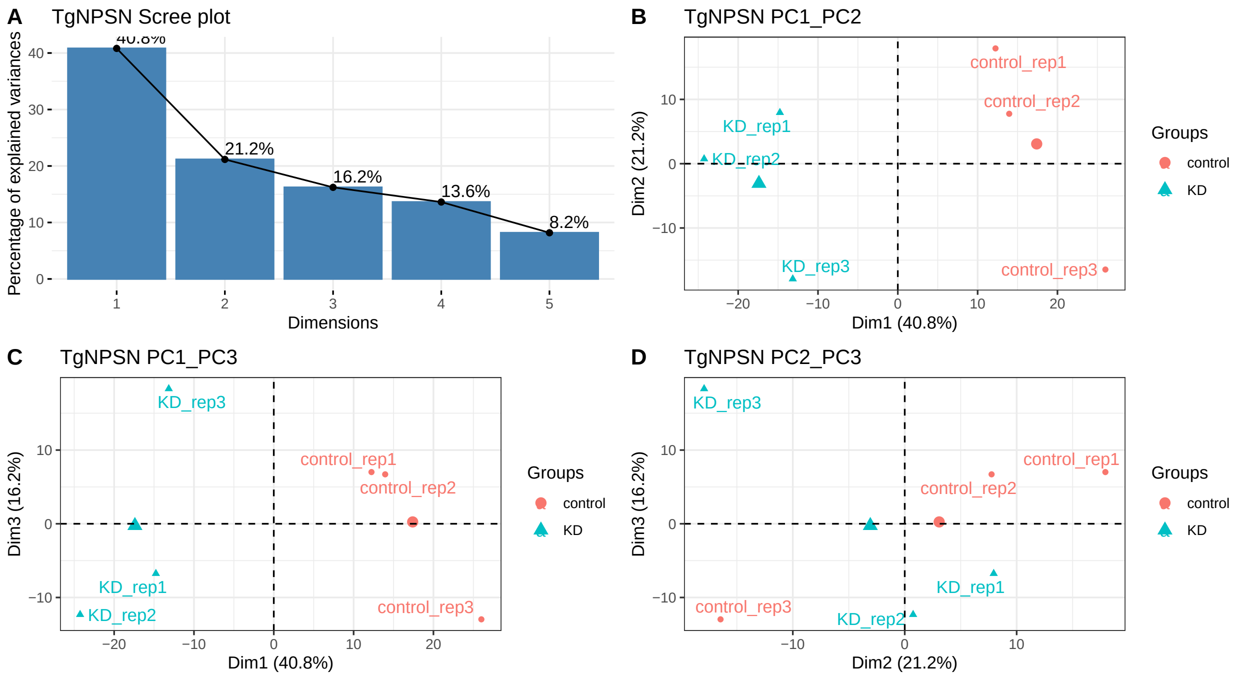

A
TgNPSN Scree plot
Percentage of explained variances
Dimensions
40.8%
21.2%
16.2%
13.6%
8.2%
B
TgNPSN PC1_PC2
Dim1 (40.8%)
Dim2 (21.2%)
control_rep1
control_rep2
control_rep3
KD_rep1
KD_rep2
KD_rep3
Groups
control
KD
C
TgNPSN PC1_PC3
Dim1 (40.8%)
Dim3 (16.2%)
D
TgNPSN PC2_PC3
Dim2 (21.2%)
Dim3 (16.2%)

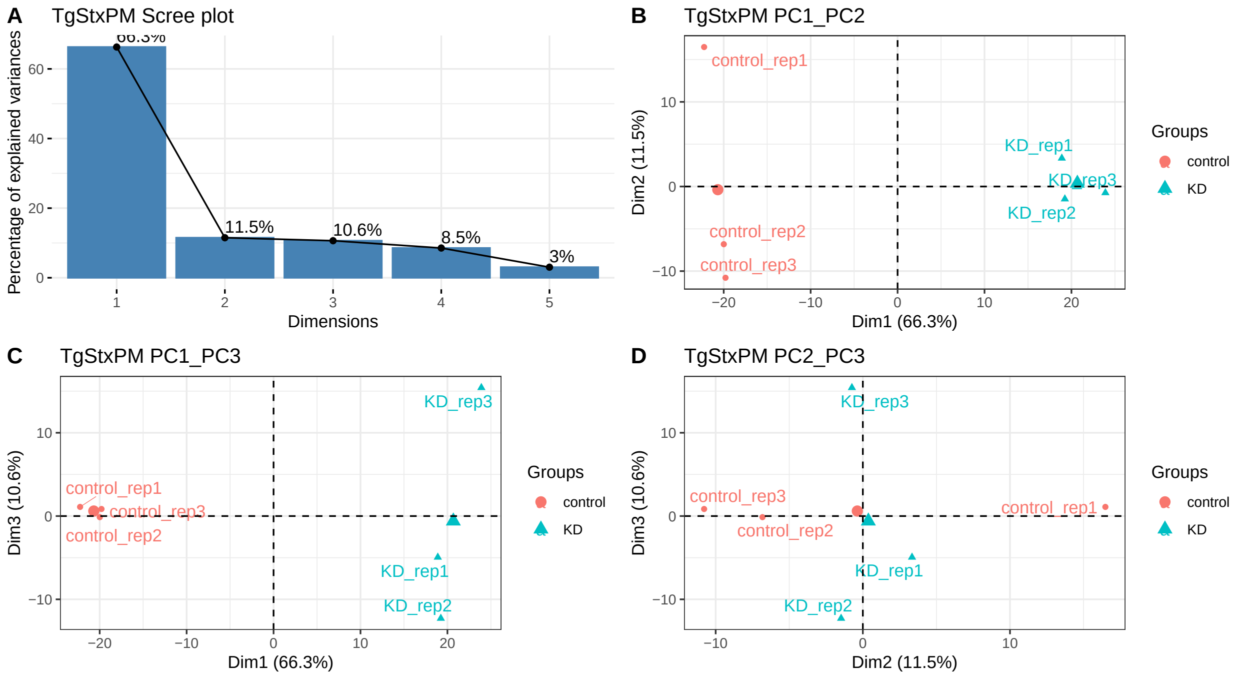


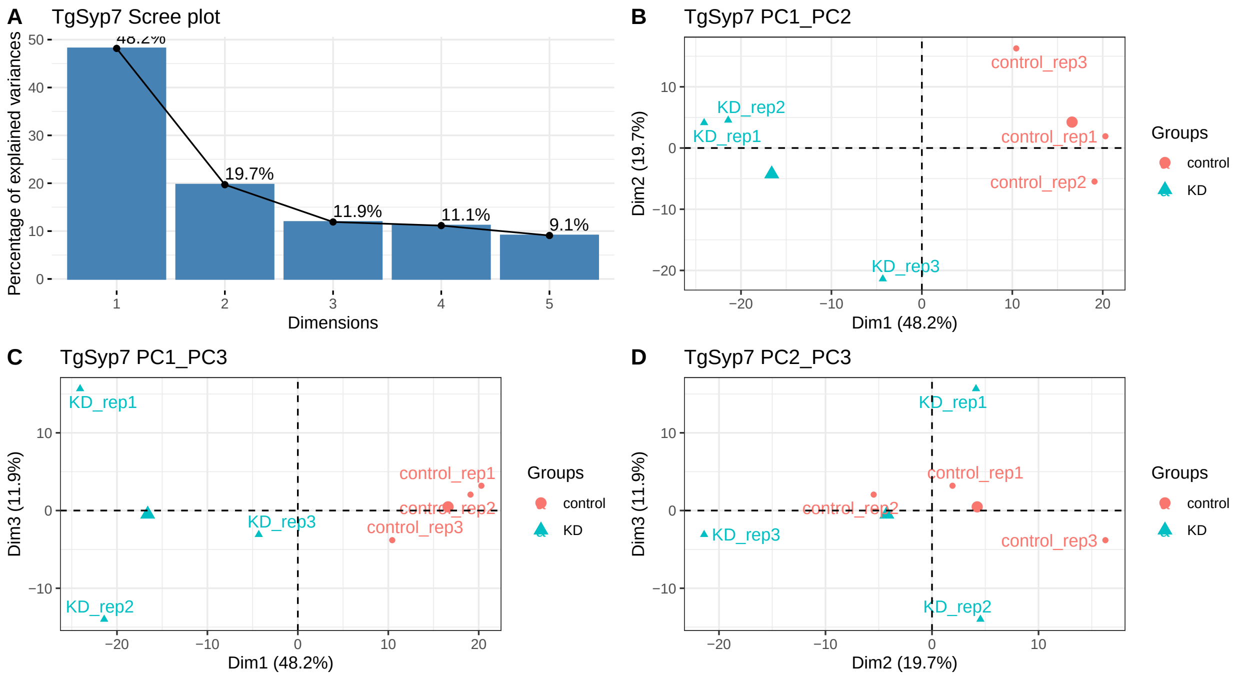


```r
#dev.off()
rm(i)
```

```r
#custom function, calculating effect size and power; input is Summarized Experiment container (obj) and
fun_power <-
  function(experiment, treat1, treat2) {

    #select protein layer from defined experiment in the list of QFeature objects
    obj <-
      ls_QFeat[[experiment]][["log2_norm_proteins_FDR5"]] %>%
```

```r
  set_colnames(colData(.)$sample.ID)

#treatment IDs
treat_ID <- c(treat1, treat2)

#wide format - each column is one replicate for treatments of interest
df <-
  assay(obj) %>%
  as.data.frame() %>%
  select(starts_with(treat_ID))

#number of replicates per treatment (assuming balanced design)
repl_n <- ncol(df)/length(treat_ID)

#calculating treatment means and SD
for (name in treat_ID) {
  df[ , paste0(name, ".mean")] <-
    df %>%
    select(starts_with(name)) %>%
    rowMeans()

  df[ , paste0(name, ".SD")] <-
    df %>%
    select(starts_with(name)) %>%
    as.matrix %>%
    rowSds()
}

#function for Cohen's D
CohenD <-
  function(obj, row, ID1, ID2, n) {
    esc_mean_sd(grp1m = obj[row, paste0(ID1, ".mean")],
                grp2m = obj[row, paste0(ID2, ".mean")],
                grp1sd = obj[row, paste0(ID1, ".SD")],
                grp2sd = obj[row, paste0(ID2, ".SD")],
                grp1n = n,
                grp2n = n)
  }


#calculating effect sizes for each entry
for (i in 1:nrow(df)) {
  esc_obj <-
    CohenD(obj = df,
           row = i,
           ID1 = treat1,
           ID2 = treat2,
           n = repl_n)

  #Hedges' g correction for small sample size (effect size becomes 0 for n=3)
  hedge_g <- hedges_g(d = esc_obj$es %>% abs(),
                      totaln = repl_n)
```

```r
      df[i , paste("CohenD", treat1, treat2, sep = ".")] <- esc_obj$es %>% abs()
      df[i , paste("HedgeG", treat1, treat2, sep = ".")] <- hedge_g
    }

    #calculating statistical power based on effect size and N for p = 0.01
    for(i in 1:nrow(df)) {
      col <-
        df %>%
        select(starts_with(paste("CohenD", treat1, treat2, sep = "."))) %>%
        colnames()

      pwr <-
        pwr.t.test(n = repl_n,
                   d = df[i, col],
                   sig.level = 0.01,
                   power = NULL,
                   type = "two.sample",
                   alternative = "two.sided")

      df[i , paste("power", treat1, treat2, sep = ".")] <- pwr$power
    }

    #column for peptide support
    pep_support <-
      obj %>% rowData() %>% as.data.frame() %>%
      select(.n) %>%
      set_colnames(value = "n.peptides") %>%
      rownames_to_column(var = "Accession")

    #joining peptide support and all calculated data
    df1 <-
      left_join(x = pep_support,
                y = df %>% rownames_to_column(var = "Accession"),
                by = "Accession")

    df1$Accession <-
      str_remove(string = df1$Accession,
                 pattern = "-t26_1-p1")


    return(df1)
  }

#applying power function across all contrasts of interest across all experiments
ls_power_dfs <-
  apply(X = contrasts, MARGIN = 1, FUN = function(x) {

    fun_power(experiment = x["experiment"],
              treat1 = x["treat1"],
              treat2 = x["treat2"])
  }) %>%
  set_names(nm_cell_lines)
```

```r
power_plots <-
  lapply(X = names(ls_power_dfs),
         FUN = function(i) {

           obj <- ls_power_dfs[[i]]

           Cohen <- colnames(obj %>% select(starts_with("CohenD")))
           power <- colnames(obj %>% select(starts_with("power")))

           A <-
             ggplot(data = obj, aes(.data[[Cohen]])) +
             geom_histogram(closed = "right") +
             stat_bin(geom = "text", aes(label = after_stat(count),
                                         vjust = -0.5)) +
             theme_bw() +
             ggtitle("effect size (Cohen's D uncorrected") +
             xlab("difference between treatment means \ndivided by pooled SD")

           B <-
             ggplot(data = obj, aes(.data[[power]])) +
             geom_histogram(binwidth = 0.1, closed = "right") +
             stat_bin(binwidth = 0.1,
                      center = 0,
                      geom = "text", aes(label = after_stat(count),
                                         vjust = -0.5)) +
             scale_x_continuous(breaks = seq(0, 1, 0.2)) +
             theme_bw() +
             ggtitle("statistical power at p = 0.01") +
             xlab("probability that test correctly \nrejects null hypothesis")


           C <-
             ggplot(obj, aes(x = .data[[Cohen]],
                             y = .data[[power]])) +
             geom_point() +
             geom_hline(yintercept = 0.8,
                        linetype = "dashed",
                        colour = "red") +
             theme_bw() +
             xlab("effect size") +
             ylab("power")

           D <-
             ggMarginal(C, type = "histogram", binwidth = 0.1, size = 10)

           plots <-
             plot_grid(A, B, D,
                       ncol = 3,
                       labels = "AUTO")

                      #add title to each plot
           title <-
             ggdraw() +
```

```r
            draw_label(label = i,
                       fontface = 'bold',
                       x = 0,
                       hjust = 0) +
            theme(plot.margin = margin(0, 0, 0, 7))

          graphs_power <-
            plot_grid(title,
                      plots,
                      ncol = 1,
                      rel_heights = c(0.1, 1))

          return(graphs_power)

        }) %>%
    set_names(nm_cell_lines)

#pdf(file = "graphs/power_plots.pdf", width = 16)
power_plots
```

```
## $TgLMBD3
```

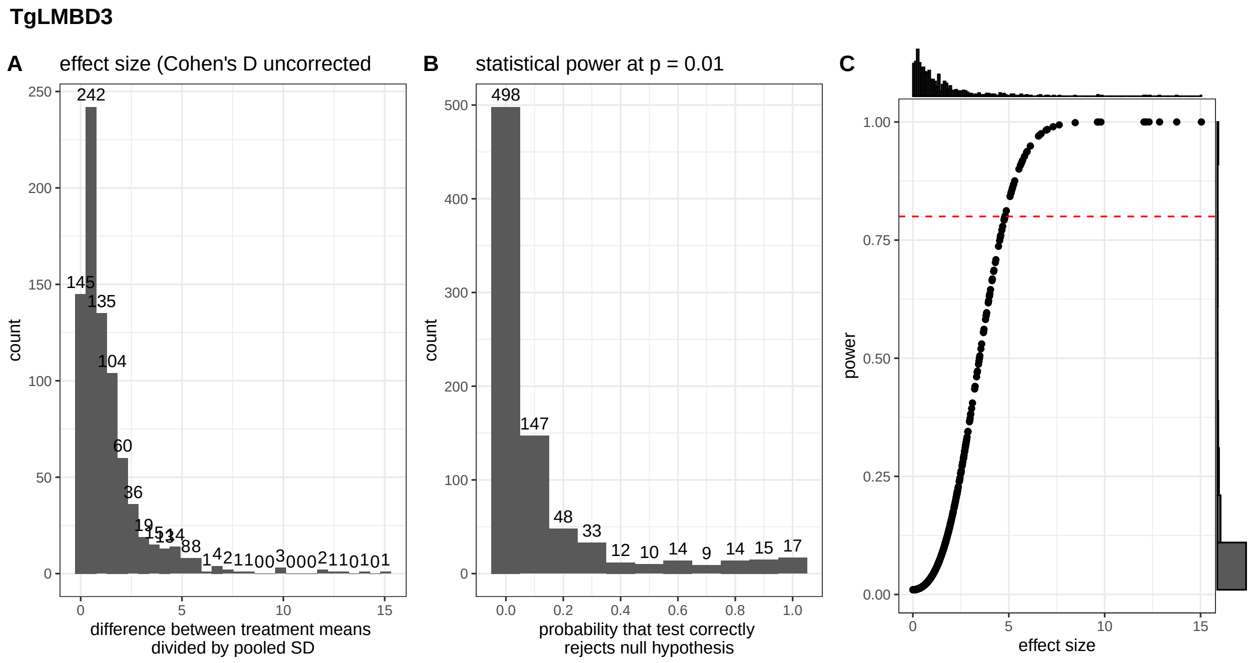


```
##
## $TgNPSN
```

**TgNPSN**

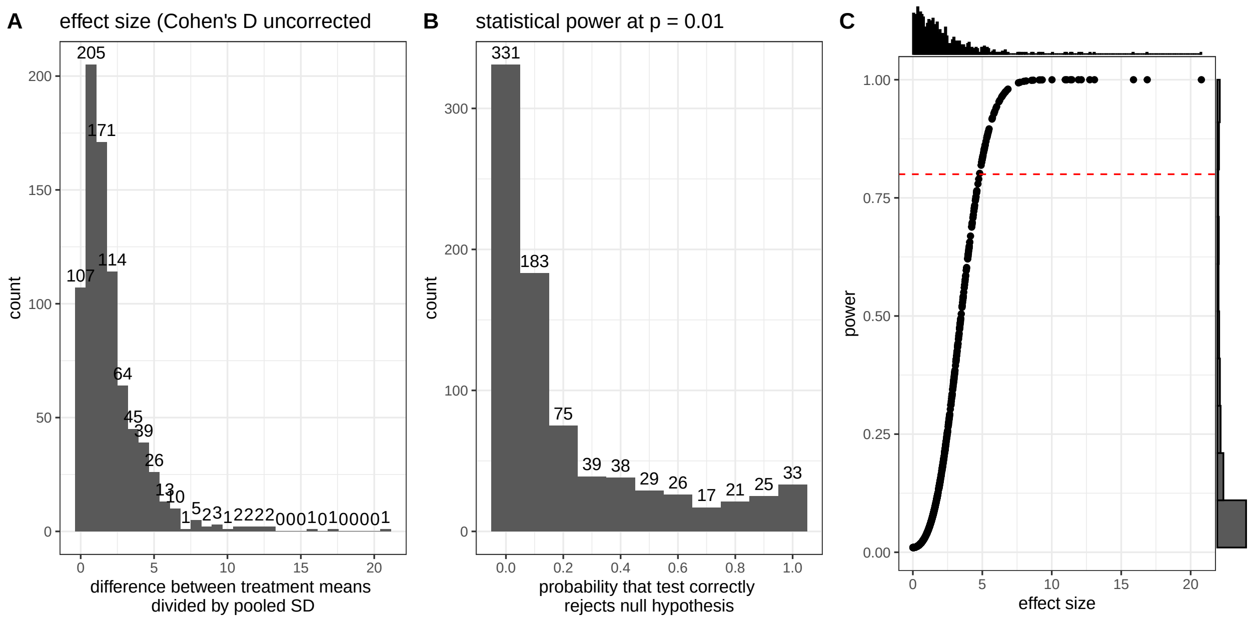


```
##
## $TgStxPM
```

**TgStxPM**

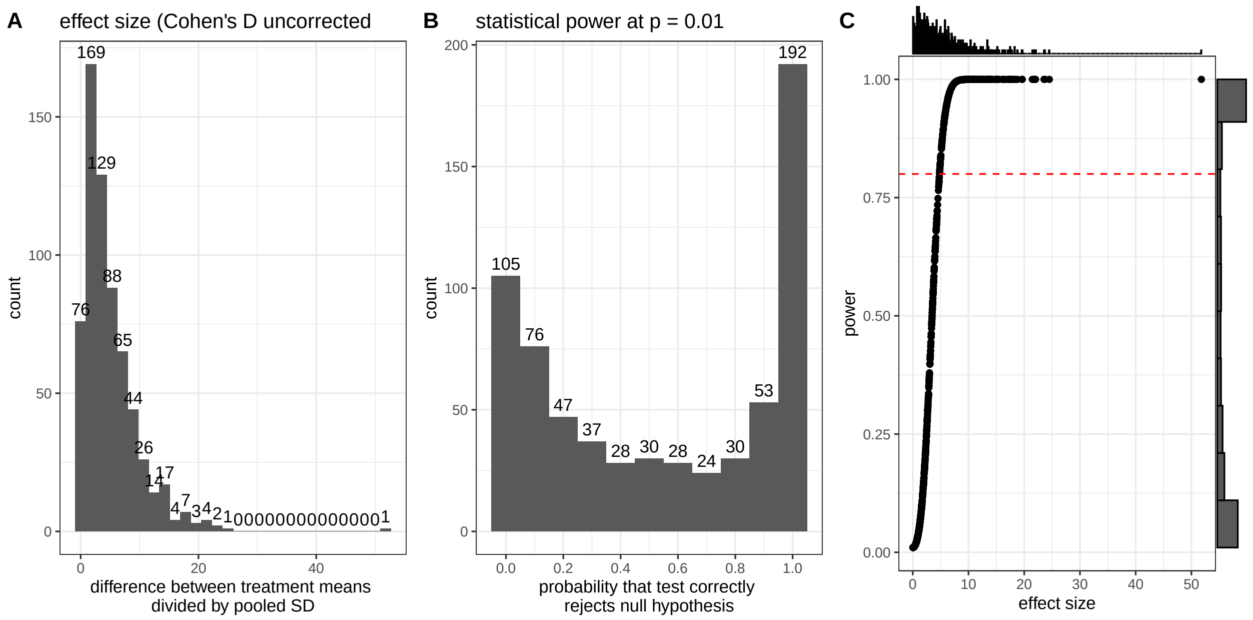


```
##
## $TgSyp7
```

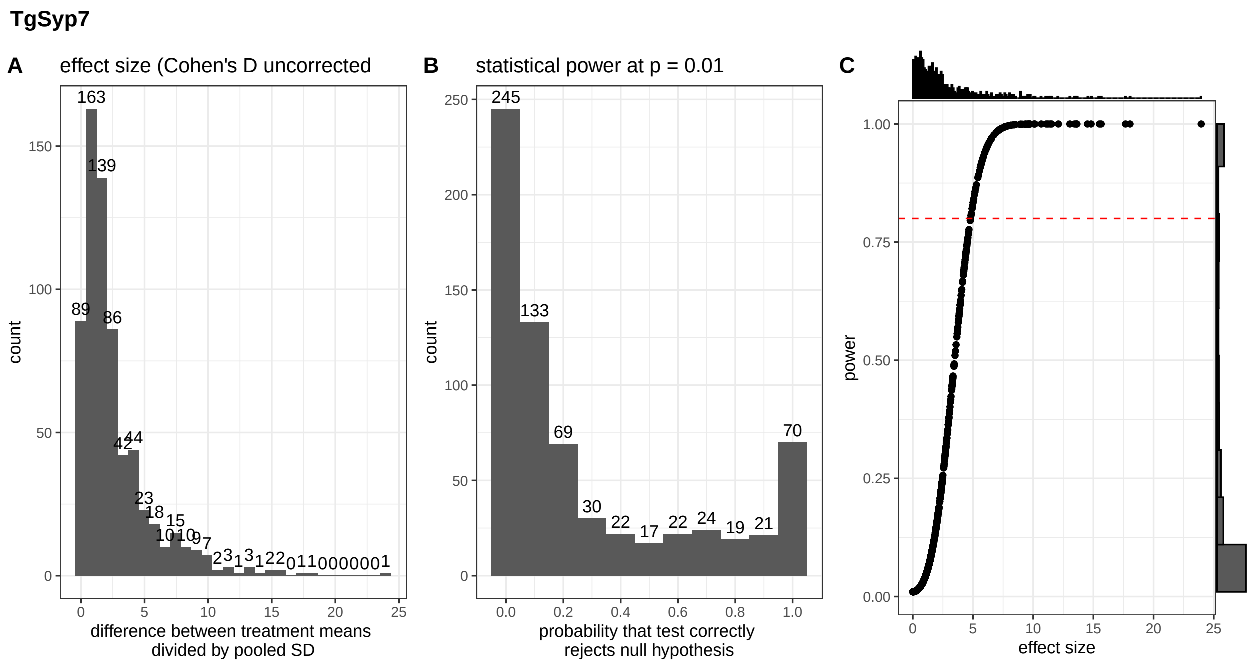


```r
#dev.off()
```

```r
ls_fit_Bayes_models <-
  lapply(X = names(ls_QFeat),
         FUN = function(i) {

           #get object
           obj <- ls_QFeat[[i]]

           #Data
           data <- obj[["log2_norm_proteins_FDR5"]] %>% assay()

           #Create model matrix
           treatment <- factor(colData(obj)$condition)
           model_design <- model.matrix(~ 0 + treatment) #a means model

           message("model design:")
           print(i)
           print(model_design)

           #define contrasts of interest
           model_contrasts <-
             makeContrasts(KD_control = treatmentKD-treatmentcontrol,
                           levels = colnames(model_design))

           message("model contrasts:")
           print(i)
           print(model_contrasts)

           #perform linear model fit
           fitted_model <- lmFit(object = data,
```

```r
                                 design = model_design)

           #Apply contrasts
           fitted_model <- contrasts.fit(fit = fitted_model,
                                         contrasts = model_contrasts)

           #Calculate test statistic using Bayes moderation of the standard errors towards a global val
           fit_Bayes <- eBayes(fitted_model,
                               trend = TRUE,
                               robust = TRUE)

           #Plot residual standard deviation vs average log abundance
           fit_Bayes %>% plotSA(xlab = "Average log2_abundance", cex = 0.5)

           return(fit_Bayes)

         }) %>%
  set_names(nm_cell_lines)
```

```
## model design:

## [1] "TgLMBD3"
##   treatmentcontrol treatmentKD
## 1                0           1
## 2                0           1
## 3                0           1
## 4                1           0
## 5                1           0
## 6                1           0
## attr(,"assign")
## [1] 1 1
## attr(,"contrasts")
## attr(,"contrasts")$treatment
## [1] "contr.treatment"

## model contrasts:

## [1] "TgLMBD3"
##                   Contrasts
## Levels              KD_control
##   treatmentcontrol         -1
##   treatmentKD               1

## model design:

## [1] "TgNPSN"
##   treatmentcontrol treatmentKD
## 1                0           1
## 2                0           1
## 3                0           1
## 4                1           0
## 5                1           0
```

```
## 6                  1           0
## attr(,"assign")
## [1] 1 1
## attr(,"contrasts")
## attr(,"contrasts")$treatment
## [1] "contr.treatment"
```

```
## model contrasts:
```

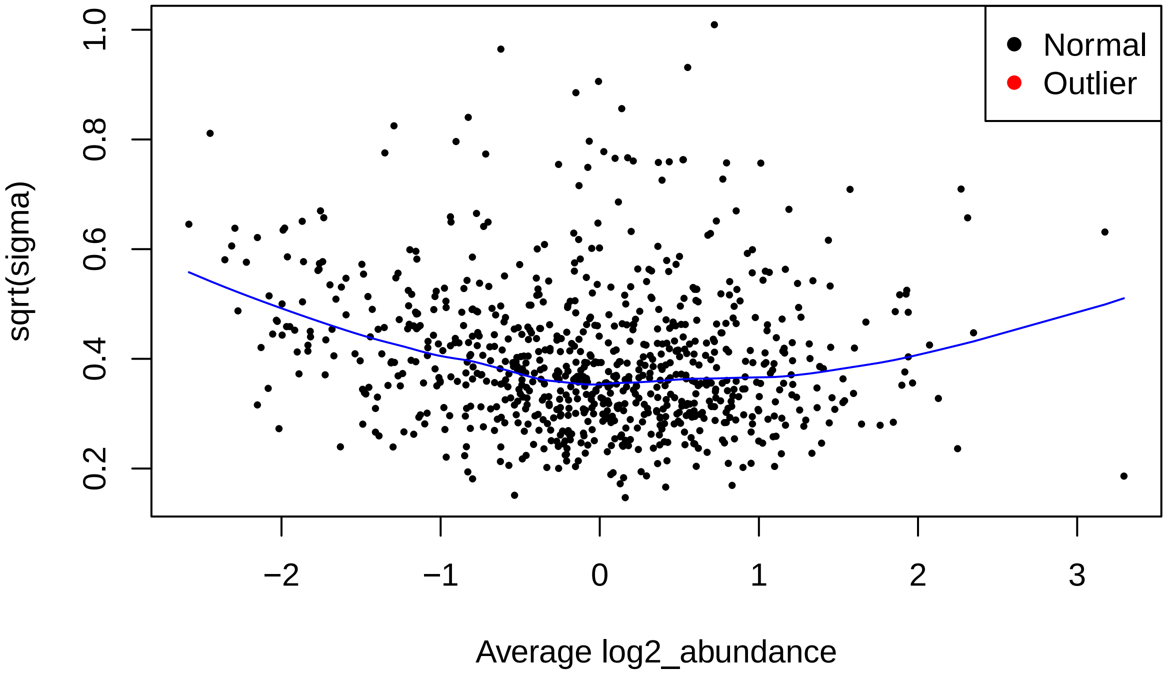


```
## [1] "TgNPSN"
##                     Contrasts
## Levels               KD_control
##   treatmentcontrol           -1
##   treatmentKD                 1
```

```
## model design:
```

```
## [1] "TgStxPM"
##   treatmentcontrol treatmentKD
## 1                0           1
## 2                0           1
## 3                0           1
## 4                1           0
## 5                1           0
## 6                1           0
```

```
## attr(,"assign")
## [1] 1 1
## attr(,"contrasts")
## attr(,"contrasts")$treatment
## [1] "contr.treatment"

## model contrasts:
```

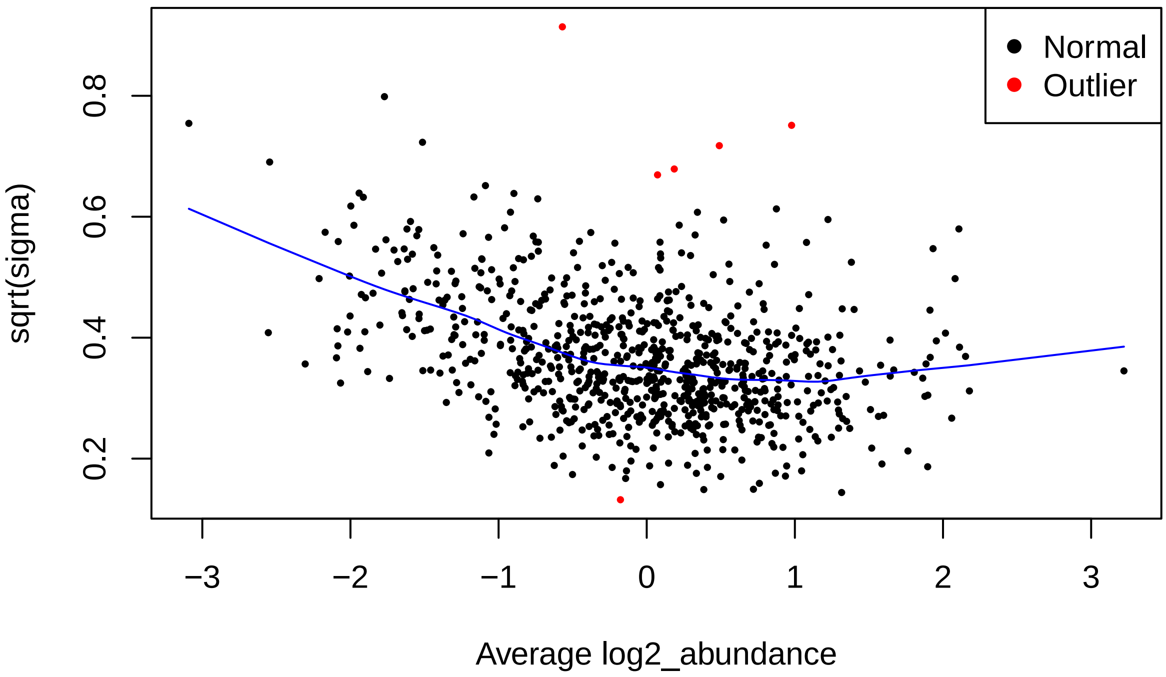


```
## [1] "TgStxPM"
##                    Contrasts
## Levels               KD_control
##   treatmentcontrol          -1
##   treatmentKD                1

## model design:

## [1] "TgSyp7"
##   treatmentcontrol treatmentKD
## 1                0           1
## 2                0           1
## 3                0           1
## 4                1           0
## 5                1           0
## 6                1           0
## attr(,"assign")
```

```
## [1] 1 1
## attr(,"contrasts")
## attr(,"contrasts")$treatment
## [1] "contr.treatment"
```

```
## model contrasts:
```

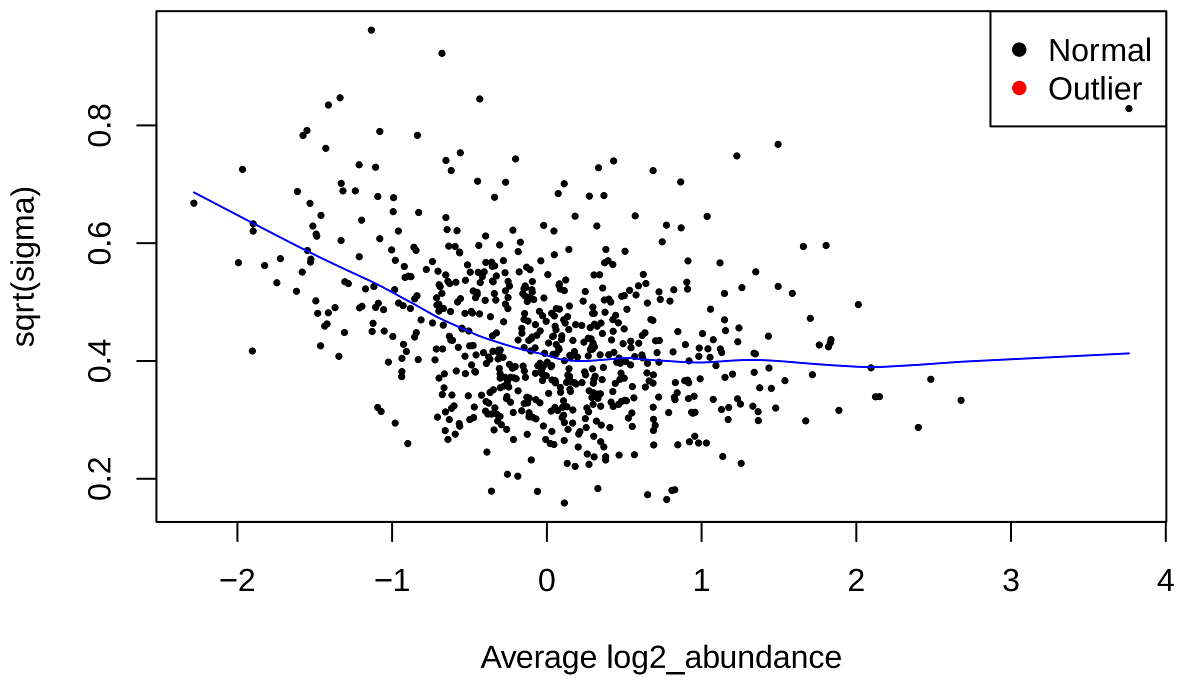


```
## [1] "TgSyp7"
##                    Contrasts
## Levels              KD_control
##   treatmentcontrol          -1
##   treatmentKD                1
```

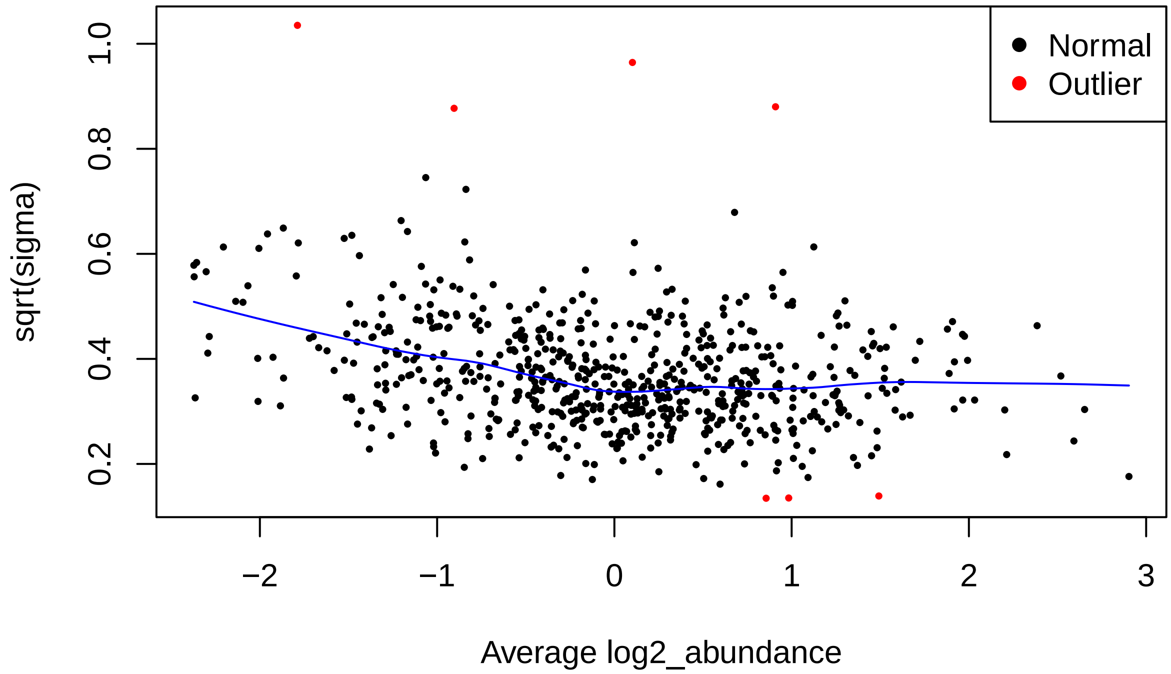


```
#HyperLOPIT annotation data from Barylu k et al. 2020 https://doi.org/10.1016/j.chom.2020.09.011 - Table
df_LOPIT <-
  read_csv(file = "input/1-s2.0-S193131282030514X-mmc5.csv") %>% as.data.frame()
```

```
## Rows: 3832 Columns: 8
## -- Column specification --------------------------------------------------------
## Delimiter: ","
## chr (5): Accession, Description, markers, tagm.map.allocation, tagm.map.allo...
## dbl (3): tagm.map.probability, tagm.map.outlier, tagm.map.loc.prob
##
## i Use `spec()` to retrieve the full column specification for this data.
## i Specify the column types or set `show_col_types = FALSE` to quiet this message.
```

```
#transcript product descriptions from ToxoDB-65_TgondiiME49
ToxoDB65_fasta <-
  readAAStringSet(filepath = "fasta files/ToxoDB-65_TgondiiME49_AnnotatedProteins.fasta",
                  format = "fasta",
                  use.names = TRUE)

Accession <- str_remove(string = names(ToxoDB65_fasta), pattern = "-t26_1-p1.*")
description <- str_match(string = names(ToxoDB65_fasta),
                         pattern = "transcript_product=(.*?) \\| location")[,2]

df_description <-
  cbind(Accession, description) %>%
```

```
  as.data.frame()

rm(Accession, description)

#join descriptions and LOPIT data
df_annot <- left_join(x = df_description,
                      y = df_LOPIT %>% select(Accession, tagm.map.allocation.pred),
                      by = "Accession")

#list of output dataframes
ls_output_df <-
  lapply(X = names(ls_fit_Bayes_models),
         FUN = function(i) {

           #get object
           obj <- ls_fit_Bayes_models[[i]]

           #output dataframe
           df <-
             topTable(obj,
                      coef = NULL,
                      adjust.method = "BH",
                      number = Inf) %>%
             rownames_to_column(var = "Accession")

           #removing "-t26_1-p1"
           df$Accession <-
             str_remove(string = df$Accession,
                        pattern = "-t26_1-p1")

           #add annotation data columns
           df2 <-
             left_join(x = df,
                       y = df_annot,
                       by = "Accession")

           return(df2)
         }) %>%
  set_names(nm_cell_lines)


#write as excel summary file
sheets <- list(TgLMBD3_abundance_data = ls_output_df[["TgLMBD3"]],
               TgNPSN_abundance_data = ls_output_df[["TgNPSN"]],
               TgSyp7_abundance_data = ls_output_df[["TgSyp7"]])

write_xlsx(x = sheets,
           path = paste0("files/summary_foldchange_data.xlsx"),
           col_names = TRUE,
           format_headers = TRUE)
```

```
#p-value distribution histograms
p_value_plots <-
  lapply(X = names(ls_output_df),
         FUN = function(i) {

           #get object
           obj <- ls_output_df[[i]]

           #plot histogram of p-value frequency
           plot <-
             obj %>%
             ggplot(aes(x = P.Value)) +
             geom_histogram(binwidth = 0.025) +
             theme_bw() +
             ggtitle(paste("Histogram of p-values for drug contrast in",
                           i,
                           "\n(Limma eBayes trend model)"))

           return(plot)

         })

#pdf(file = "graphs/p-value_distribution.pdf")
for (i in seq_along(p_value_plots)) {
  plot(p_value_plots[[i]])
      }
```

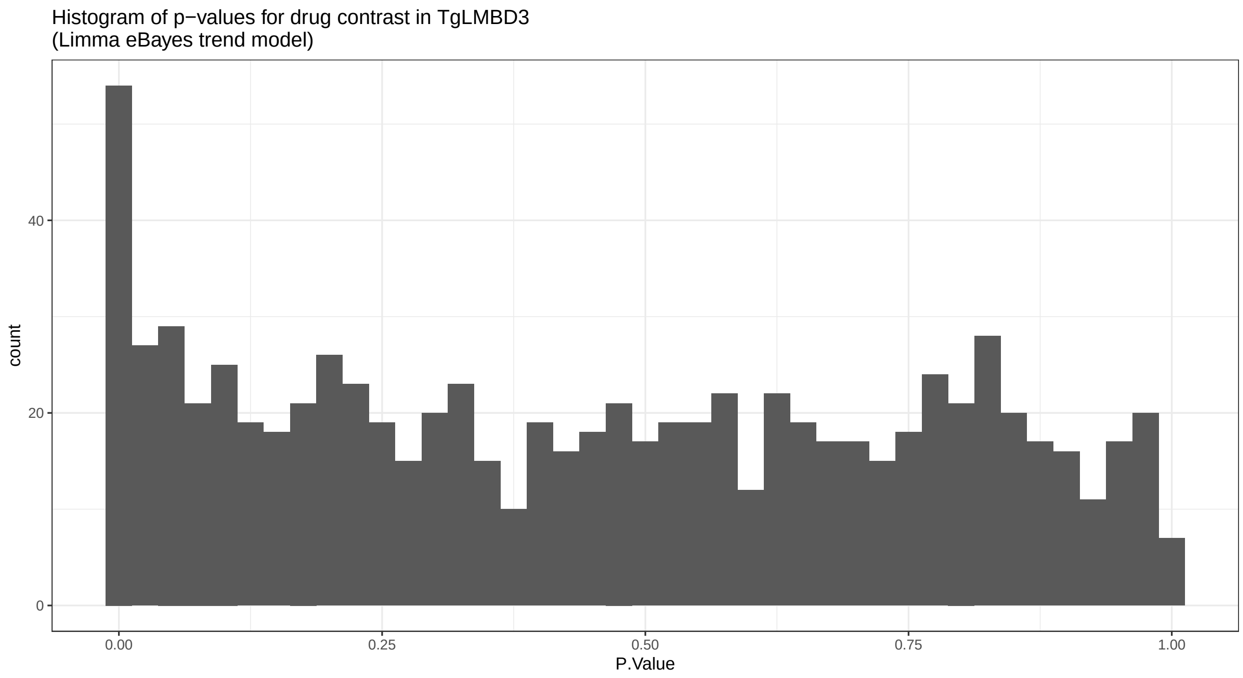

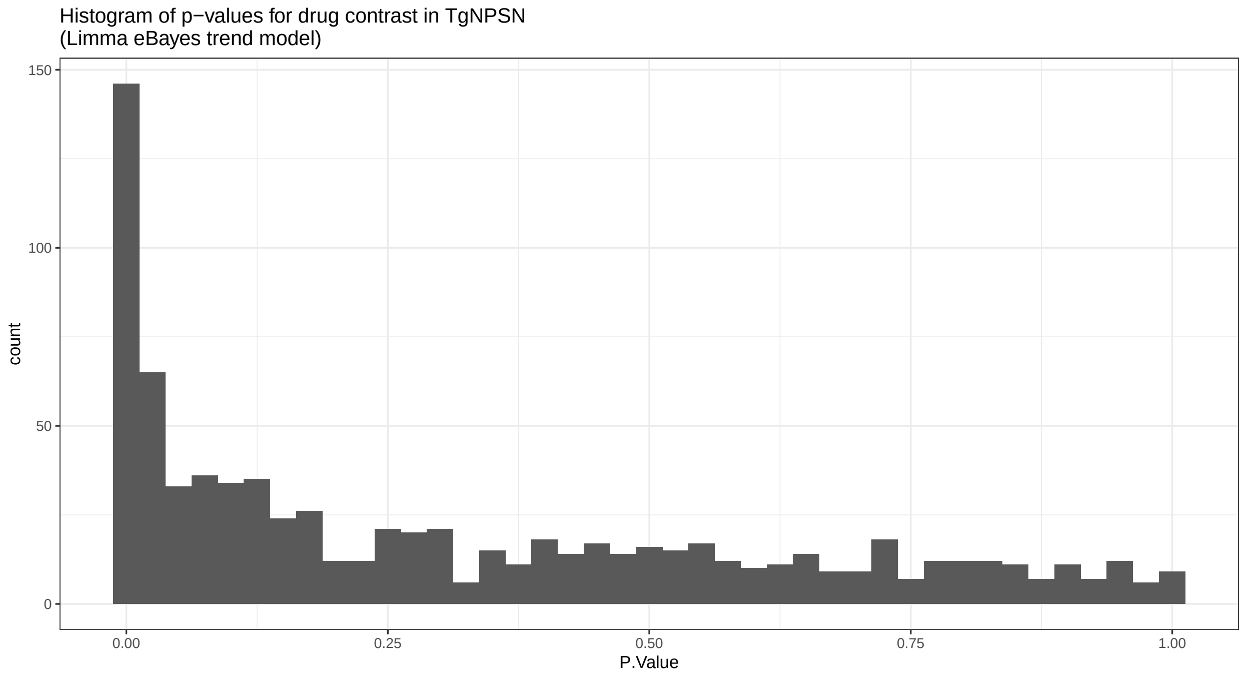
Histogram of p−values for drug contrast in TgNPSN
(Limma eBayes trend model)
count
150
100
50
0
0.00
0.25
0.50
0.75
1.00
P.Value

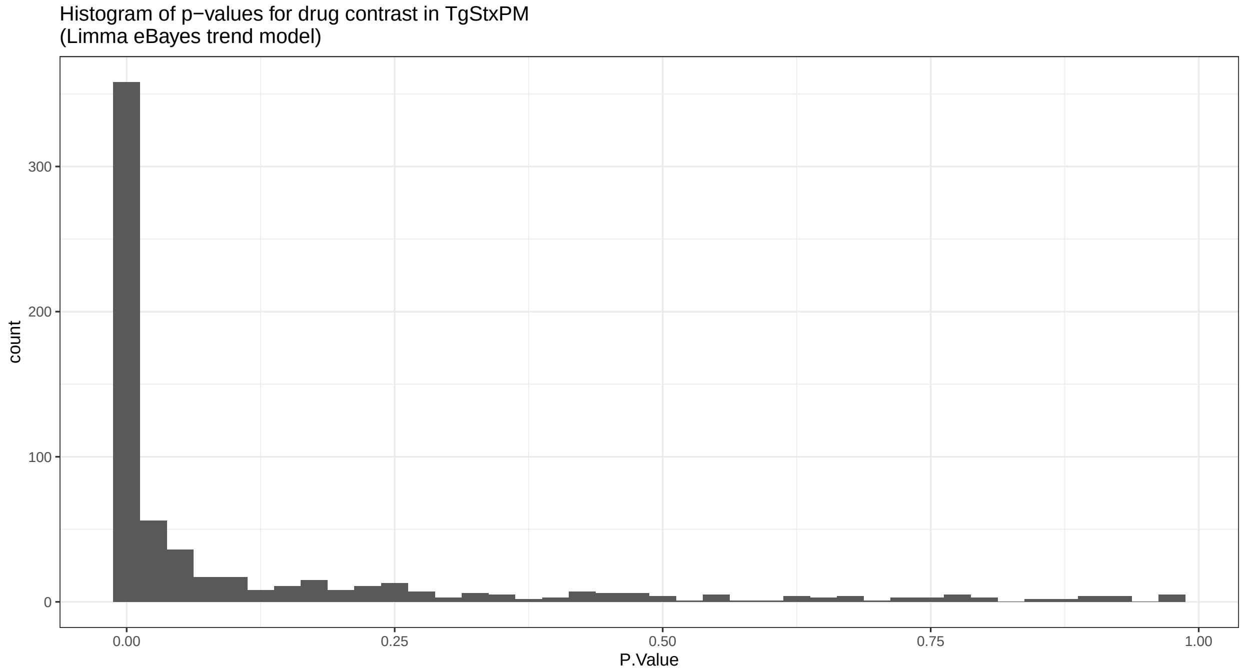
Histogram of p−values for drug contrast in TgStxPM
(Limma eBayes trend model)
count
300
200
100
0
0.00
0.25
0.50
0.75
1.00
P.Value

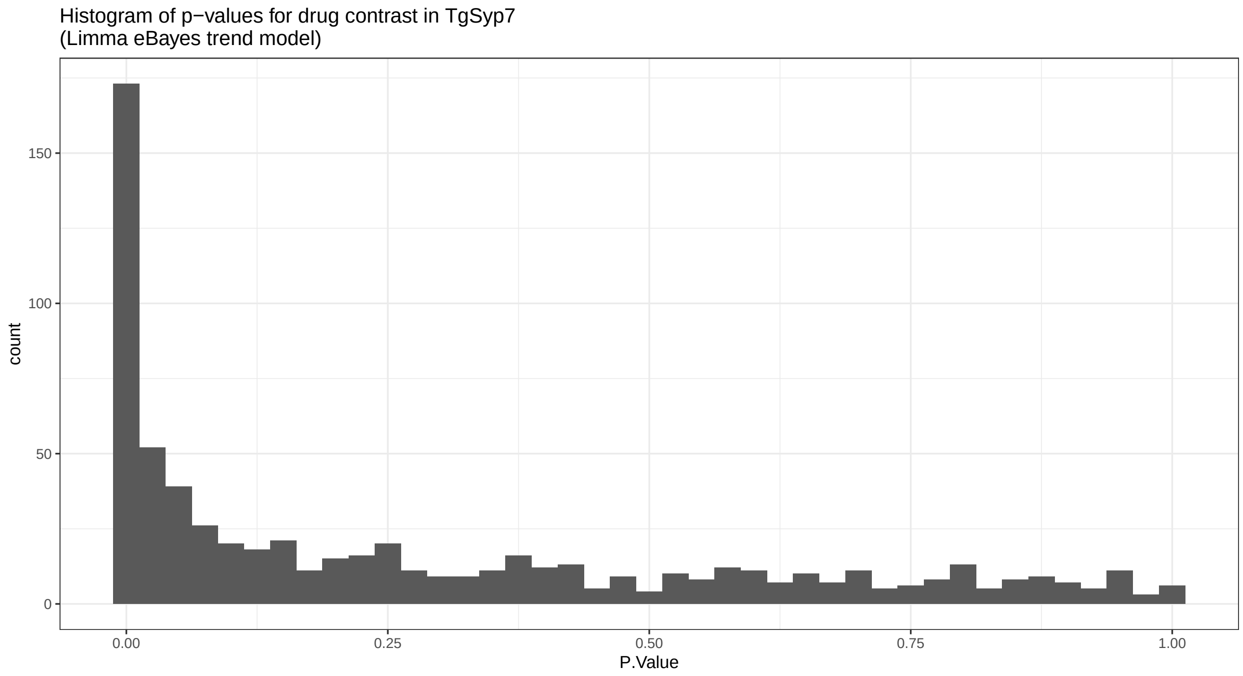


```
#dev.off()
```

```
#creating empty list
ls_all_data <-
  vector(mode = "list",
         length = length(nm_cell_lines)) %>%
  set_names(nm_cell_lines)

#filling empty list with joined dataframes from power and contrast data
for(i in 1:length(nm_cell_lines)) {

  ls_all_data[[i]] <-
    left_join(x = ls_power_dfs[[i]],
              y = ls_output_df[[i]],
              by = "Accession")
  }

invisible(
  lapply(X = names(ls_all_data),
         FUN = function(i) {

           obj <- ls_all_data[[i]]

           write_csv(obj,
                     file = paste0("files/",
                                   i,
                                   "_all_data.csv"))
         })
)
```

```r
#Volcano plot with conditional colouring
ls_plots <-
  lapply(X = names(ls_all_data),
         FUN = function(i){

           obj <- ls_all_data[[i]]

           #add colour column based on p.adj and log2FC for Volcano plot
           obj <-
             obj %>%
             mutate(Volcano.colour = case_when(tagm.map.allocation.pred == "dense granules" &
                                                 adj.P.Val <= 0.05 ~ "red",
                                               tagm.map.allocation.pred == "dense granules" &
                                                 adj.P.Val > 0.05 ~ "orange",
                                               TRUE ~ "black"))

           plot <-
             ggplot(data = obj %>% arrange(Volcano.colour),
                    mapping = aes(x = logFC,
                                  y = P.Value,
                                  colour = Volcano.colour)) +
             geom_point() +
             theme_bw() +
             scale_y_continuous(trans = compose_trans("log10", "reverse")) +
             scale_color_identity() +
             xlab(label = "log2-fold change") +
             ylab(label = "p-value") +
             geom_vline(xintercept = 0, linetype = "dashed") +
             ggtitle(i)


           return(plot)

         }) %>%
  set_names(nm_cell_lines)

#save all Volcano plots in individual experiment pdfs
#pdf(file = "graphs/Volcano_plots_significance.pdf")
invisible(
  lapply(X = names(ls_plots),
                 FUN = function(i) {

                   plot(ls_plots[[i]])

                 })
  )
```

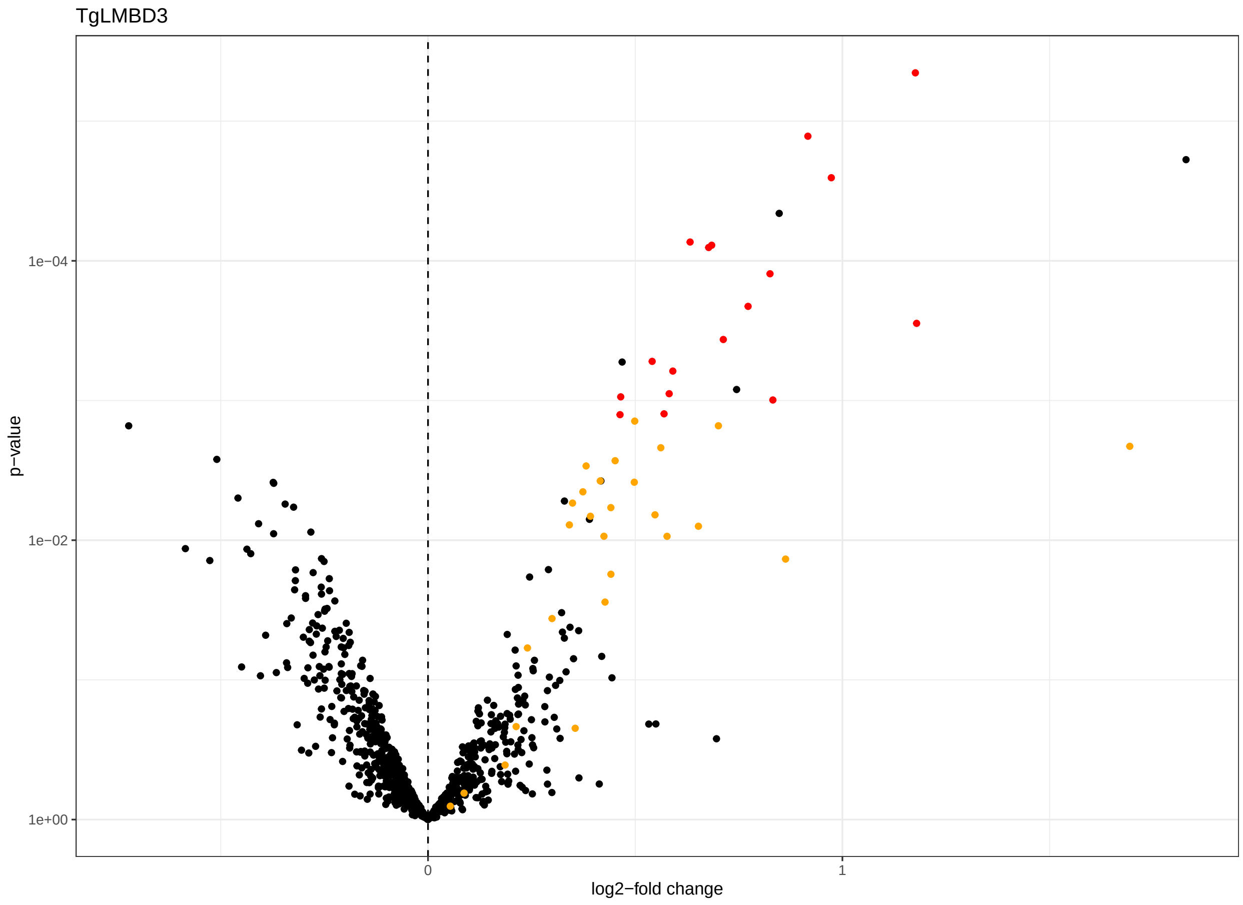

TgLMBD3
p−value
1e−04
1e−02
1e+00
0
1
log2−fold change

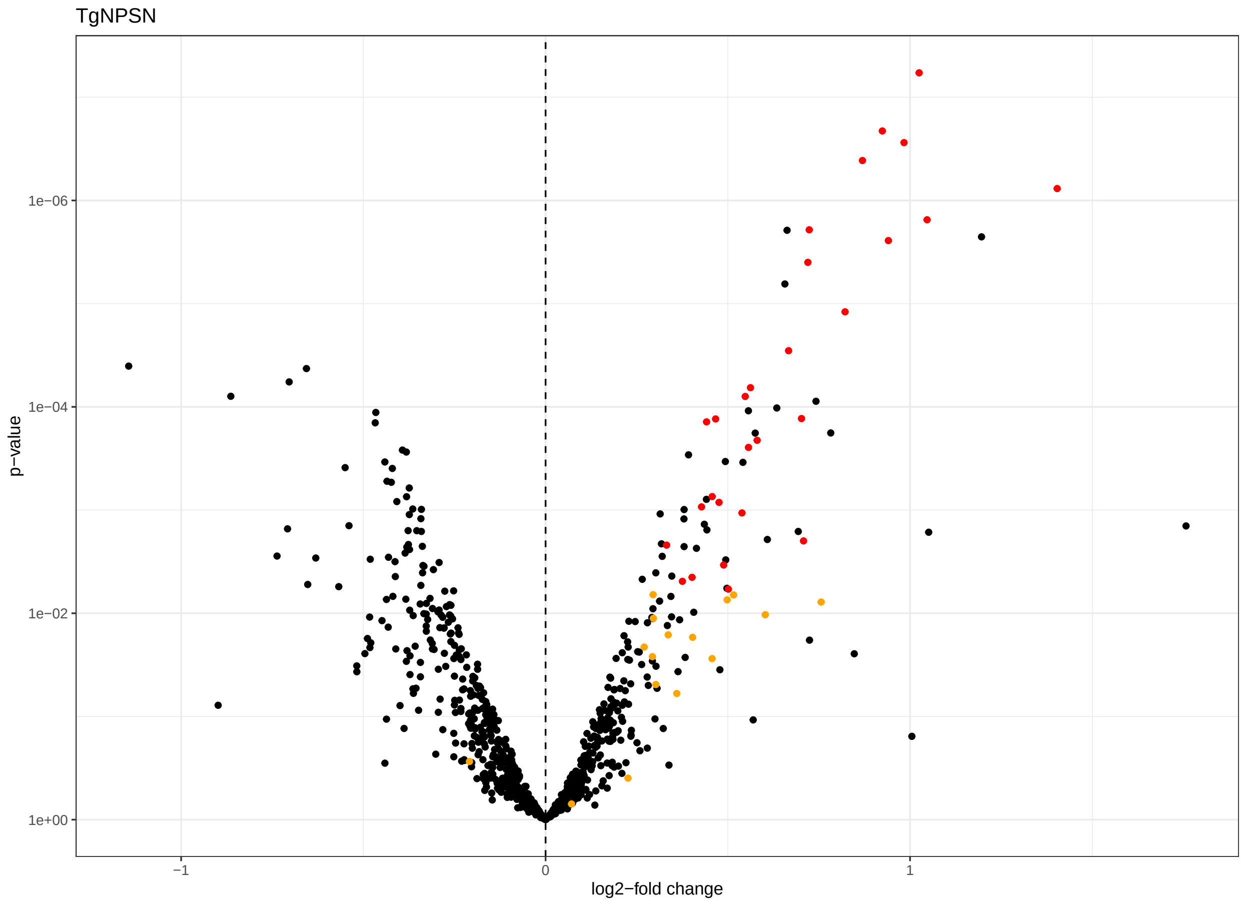
TgNPSN
p−value
1e−06
1e−04
1e−02
1e+00
−1
0
1
log2−fold change

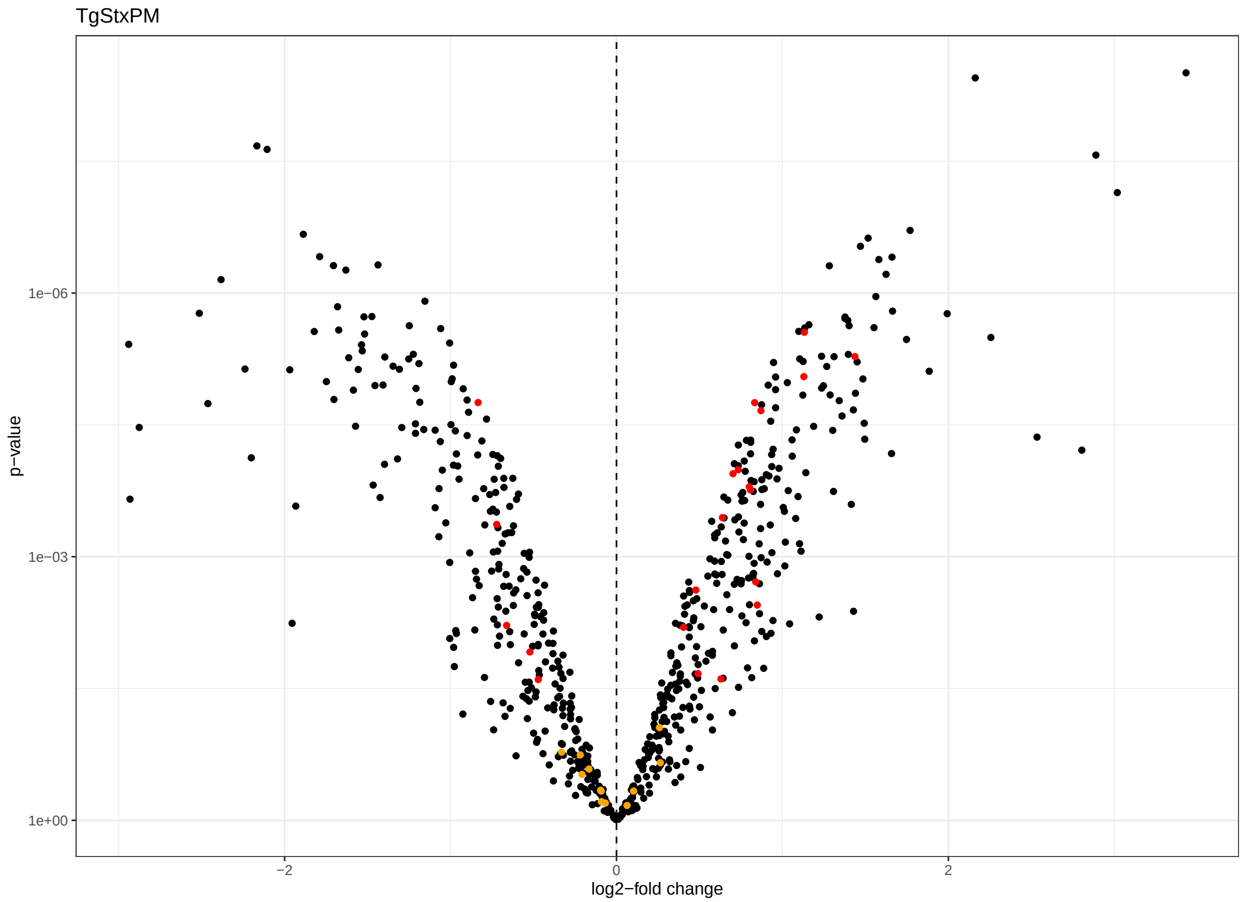
TgStxPM
p−value
1e−06
1e−03
1e+00
−2
0
2
log2−fold change

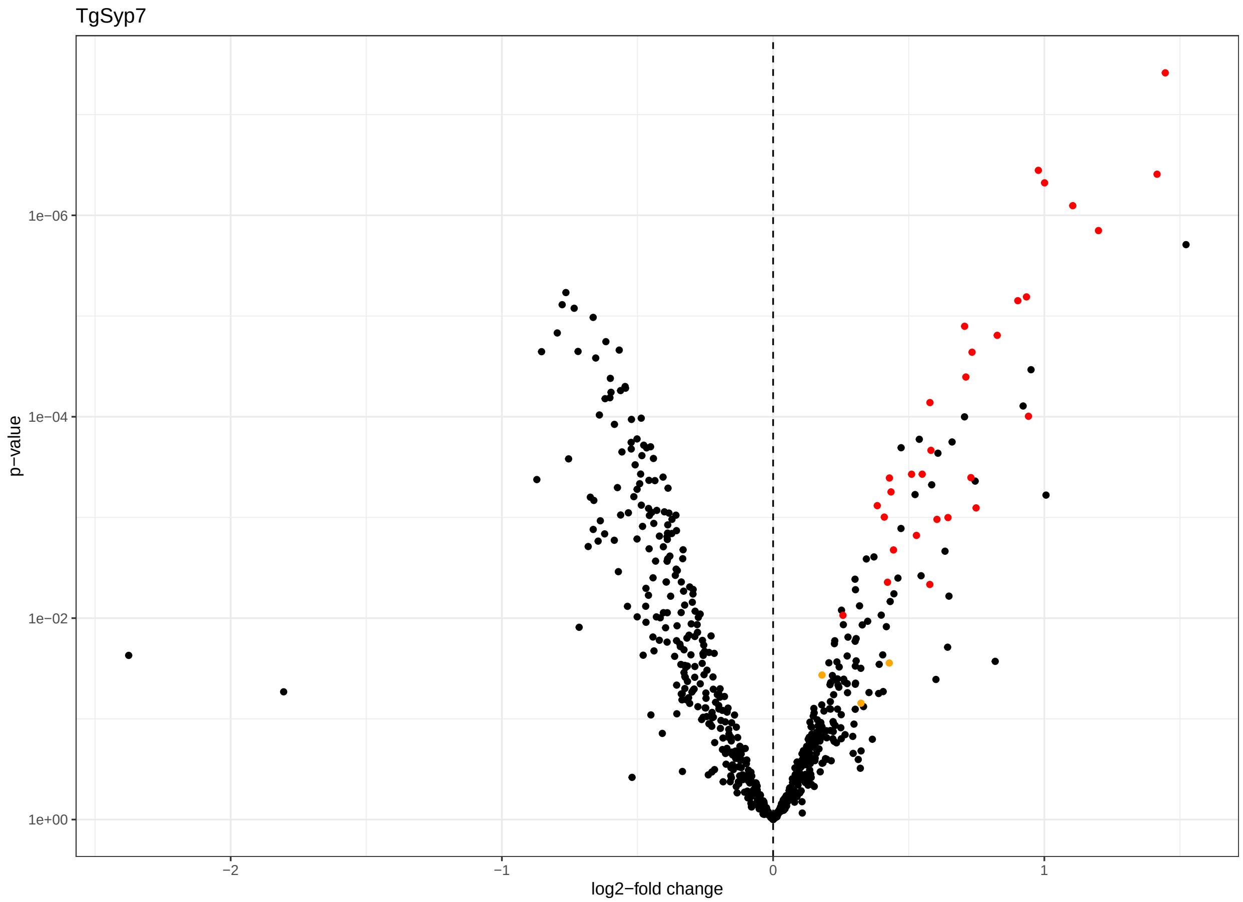


```r
#dev.off()

#save as pdf in form of a 1x3 grid
#pdf(file = "graphs/Volcano_plots_cell_lines.pdf", width = 12, height = 4)
plot_grid(plotlist = ls_plots[c("TgLMBD3",
                                "TgNPSN",
                                "TgSyp7")],
          ncol = 3,
          align = "hv",
          axis = "bl")
```

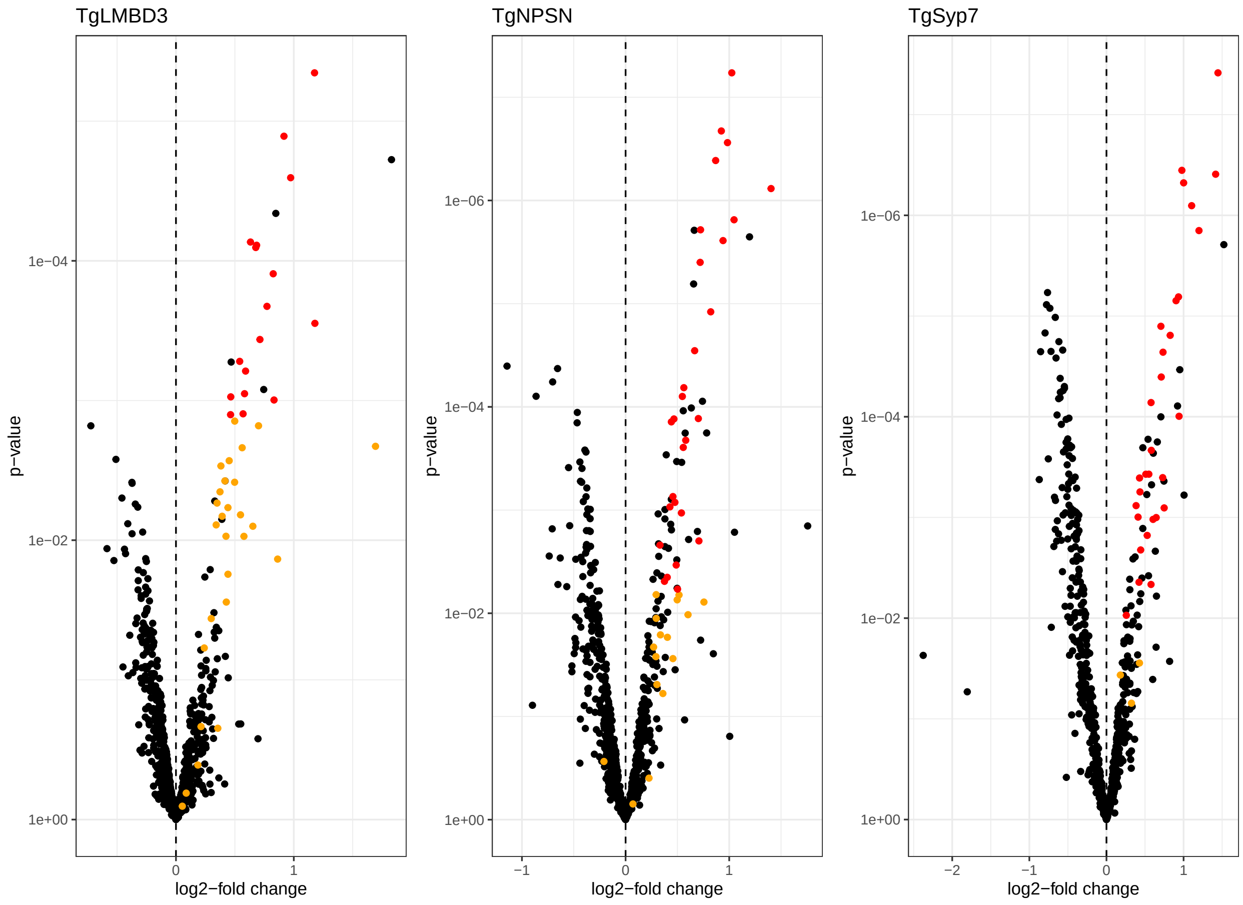


```
#dev.off()
```

```
#list of Volcano plots with power overlay
ls_plots_power <-
  lapply(X = names(ls_all_data),
         FUN = function(i) {

           #get object
           obj <- ls_all_data[[i]]

           #add colour column based on power for Volcano plot
           df <-
             obj %>%
             mutate(power.colour =
                      case_when(power.KD.control < 0.8 &
                                  adj.P.Val < 0.01 &
                                  tagm.map.allocation.pred == "dense granules" ~ "red",
                                power.KD.control >= 0.8 &
                                  adj.P.Val <= 0.01 &
                                  tagm.map.allocation.pred == "dense granules" ~ "blue",
                                TRUE ~ "black"))


           plot <-
             ggplot(data = df,
```

```
                    mapping = aes(x = logFC,  #log2FC
                                  y = P.Value,  #p.value
                                  colour = power.colour)) + #p.value.adj in colour
             geom_point() +
             theme_bw() +
             scale_y_continuous(trans = compose_trans("log10", "reverse")) +
             scale_x_continuous(breaks = seq(-6, 6, 1)) +
             #scale_colour_gradientn(colours = rainbow(7)) +
             scale_color_identity() +
             xlab(label = "log2-fold change") +
             ylab(label = "p-value") +
             geom_vline(xintercept = 0, linetype = "dashed") +
             ggtitle(i)

           return(plot)
         }) %>%
  set_names(nm_cell_lines)

#save all Volcano plots in individual experiment pdfs
#pdf(file = "graphs/Volcano_plots_power.pdf")
invisible(
  lapply(X = names(ls_plots_power),
         FUN = function(i) {

           plot(ls_plots_power[[i]])

         })
)
```

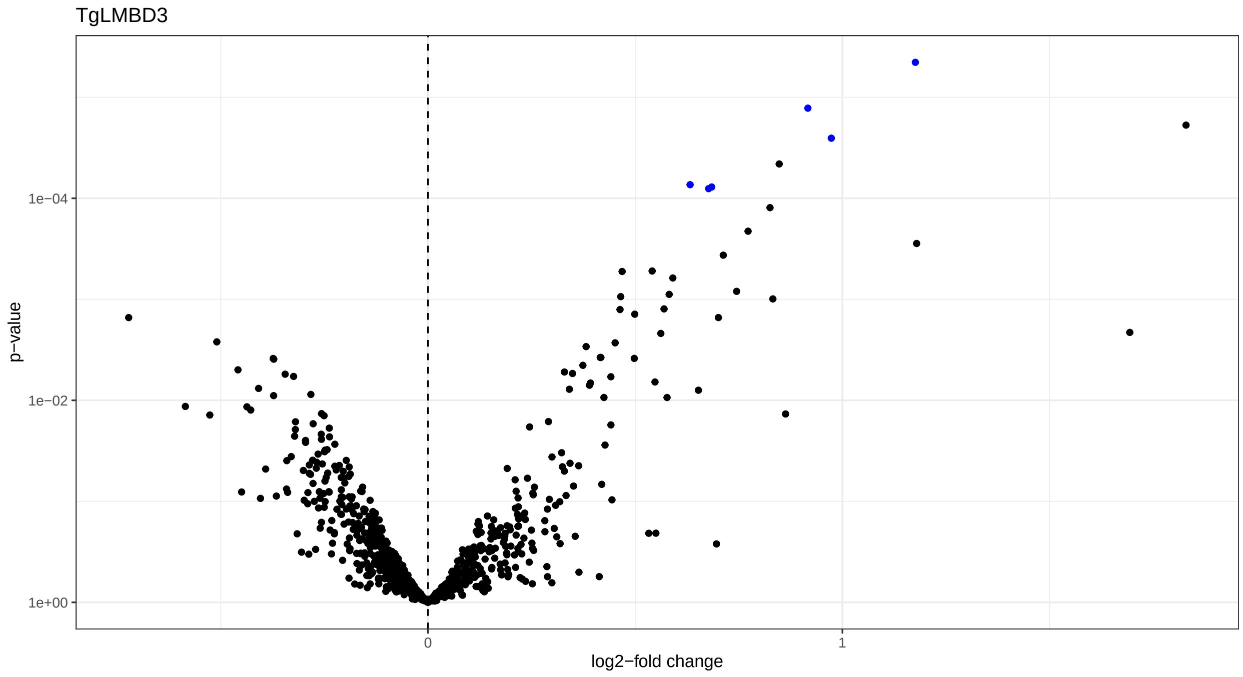

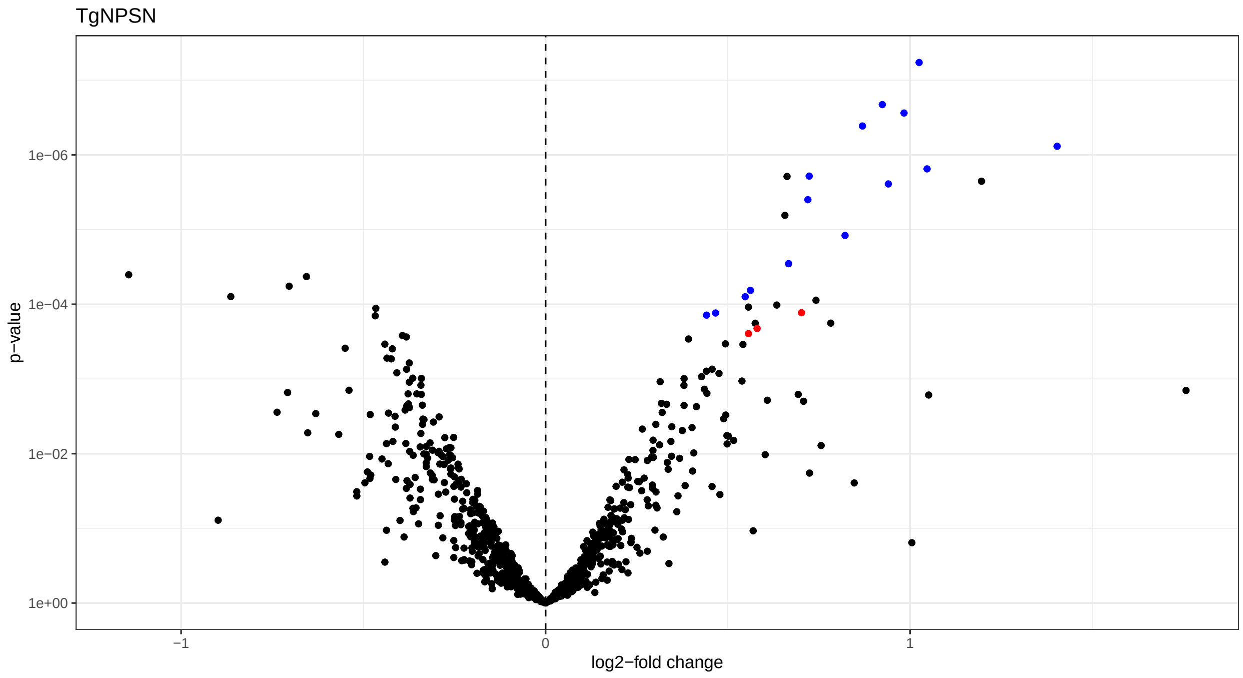
TgNPSN
p−value
1e−06
1e−04
1e−02
1e+00
−1
0
1
log2−fold change

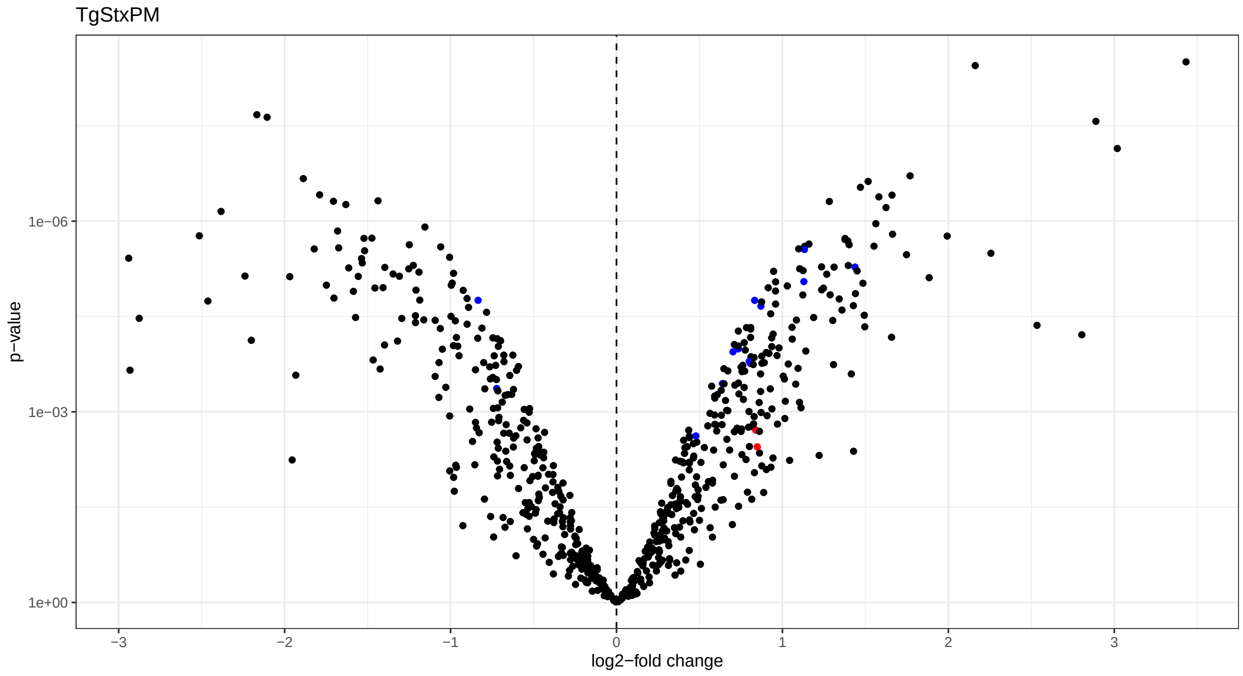
TgStxPM
p−value
1e−06
1e−03
1e+00
−3
−2
−1
0
1
2
3
log2−fold change

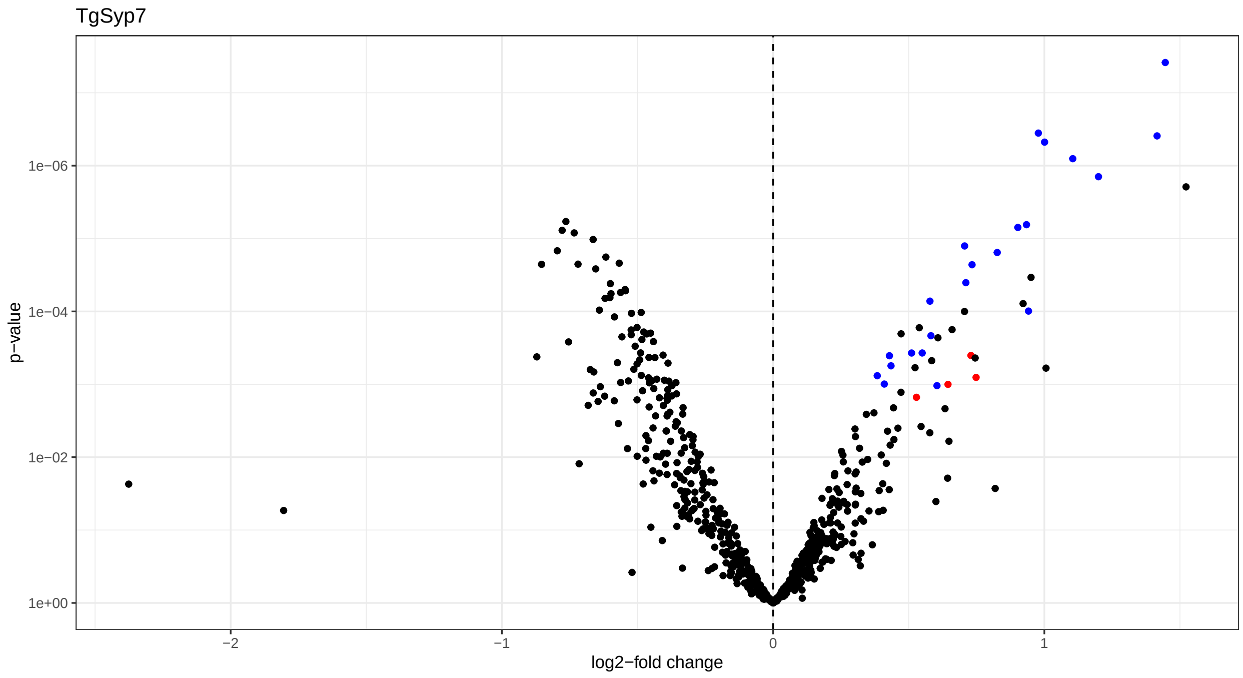


```r
#dev.off()

#save as pdf in form of a 1x3 grid
#pdf(file = "graphs/Volcano_plots_cell_lines_power.pdf", width = 12, height = 4)
plot_grid(plotlist = ls_plots_power[c("TgLMBD3",
                                      "TgNPSN",
                                      "TgSyp7")],
          ncol = 3,
          align = "hv",
          axis = "bl")
```

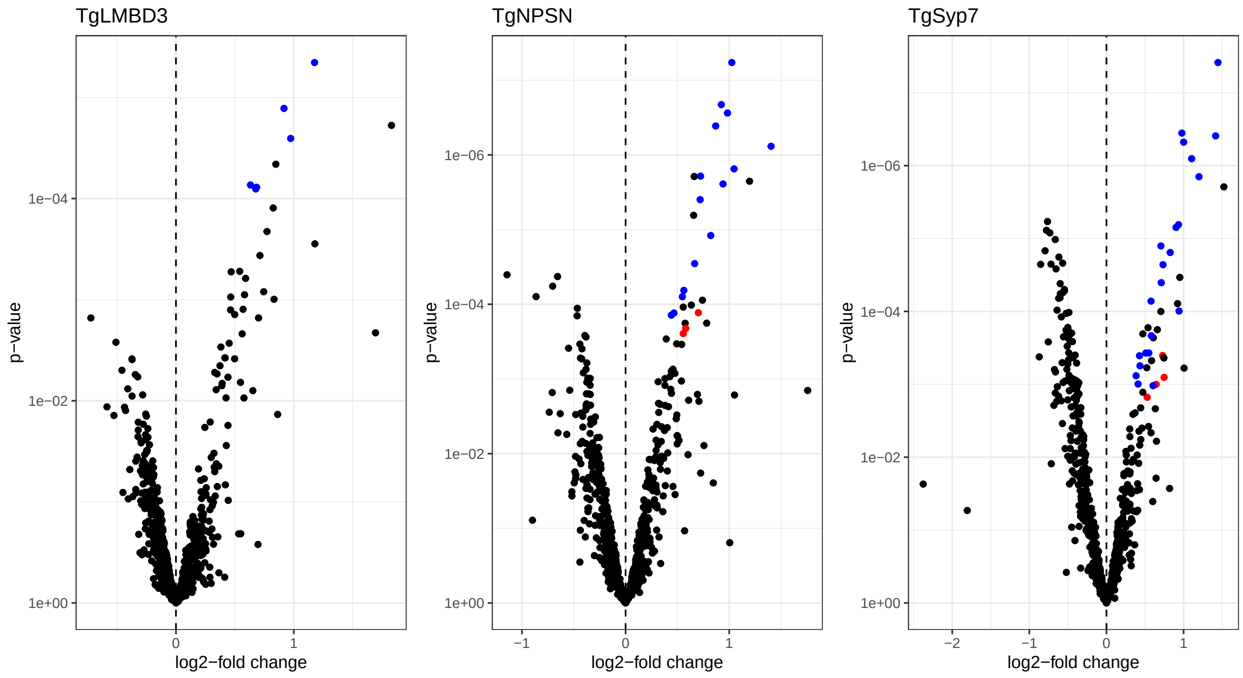


```r
#dev.off()
```

```r
sessionInfo()
```

```
## R version 4.3.1 (2023-06-16 ucrt)
## Platform: x86_64-w64-mingw32/x64 (64-bit)
## Running under: Windows 10 x64 (build 19045)
##
## Matrix products: default
##
##
## locale:
## [1] LC_COLLATE=English_United Kingdom.utf8
## [2] LC_CTYPE=English_United Kingdom.utf8
## [3] LC_MONETARY=English_United Kingdom.utf8
## [4] LC_NUMERIC=C
## [5] LC_TIME=English_United Kingdom.utf8
##
## time zone: Europe/London
## tzcode source: internal
##
## attached base packages:
## [1] stats4    stats     graphics  grDevices utils     datasets  methods
## [8] base
##
## other attached packages:
##  [1] conflicted_1.2.0           writexl_1.4.2
##  [3] Biostrings_2.68.1          XVector_0.40.0
##  [5] esc_0.5.1                  pwr_1.3-0
##  [7] ggExtra_0.10.1             ggforce_0.4.1
##  [9] limma_3.56.2               factoextra_1.0.7
```

```
## [11] FactoMineR_2.8              eulerr_7.0.0
## [13] magrittr_2.0.3              NormalyzerDE_1.18.1
## [15] QFeatures_1.10.0            MultiAssayExperiment_1.26.0
## [17] SummarizedExperiment_1.30.2 Biobase_2.60.0
## [19] GenomicRanges_1.52.0        GenomeInfoDb_1.36.3
## [21] IRanges_2.34.1              S4Vectors_0.38.1
## [23] BiocGenerics_0.46.0         MatrixGenerics_1.12.3
## [25] matrixStats_1.0.0           cowplot_1.1.1
## [27] scales_1.2.1                lubridate_1.9.2
## [29] forcats_1.0.0               stringr_1.5.0
## [31] dplyr_1.1.3                 purrr_1.0.2
## [33] readr_2.1.4                 tidyr_1.3.0
## [35] tibble_3.2.1                ggplot2_3.4.3
## [37] tidyverse_2.0.0
##
## loaded via a namespace (and not attached):
##   [1] splines_4.3.1           later_1.3.1             bitops_1.0-7
##   [4] cellranger_1.1.0        polyclip_1.10-4         preprocessCore_1.62.1
##   [7] rpart_4.1.19            lifecycle_1.0.3         rstatix_0.7.2
##  [10] lattice_0.21-8          vroom_1.6.3             MASS_7.3-60
##  [13] flashClust_1.01-2       backports_1.4.1         Hmisc_5.1-1
##  [16] rmarkdown_2.25          yaml_2.3.7              httpuv_1.6.11
##  [19] sp_2.0-0                MsCoreUtils_1.12.0      RColorBrewer_1.1-3
##  [22] abind_1.4-5             zlibbioc_1.46.0         AnnotationFilter_1.24.0
##  [25] RCurl_1.98-1.12         nnet_7.3-19             tweenr_2.0.2
##  [28] sandwich_3.0-2          GenomeInfoDbData_1.2.10 ggrepel_0.9.3
##  [31] terra_1.7-46            nortest_1.0-4           codetools_0.2-19
##  [34] DelayedArray_0.26.7     DT_0.29                 tidyselect_1.2.0
##  [37] RcmdrMisc_2.9-0         raster_3.6-23           farver_2.1.1
##  [40] base64enc_0.1-3         e1071_1.7-13            ellipsis_0.3.2
##  [43] Formula_1.2-5           emmeans_1.8.8           polylabelr_0.2.0
##  [46] tools_4.3.1             Rcpp_1.0.11             glue_1.6.2
##  [49] gridExtra_2.3           BiocBaseUtils_1.2.0     xfun_0.40
##  [52] mgcv_1.9-0              withr_2.5.1             BiocManager_1.30.22
##  [55] fastmap_1.1.1           fansi_1.0.4             digest_0.6.33
##  [58] timechange_0.2.0        R6_2.5.1                mime_0.12
##  [61] estimability_1.4.1      colorspace_2.1-0        utf8_1.2.3
##  [64] generics_0.1.3          hexbin_1.28.3           data.table_1.14.8
##  [67] class_7.3-22            htmlwidgets_1.6.2       S4Arrays_1.0.6
##  [70] scatterplot3d_0.3-44    pkgconfig_2.0.3         gtable_0.3.4
##  [73] htmltools_0.5.6         carData_3.0-5           multcompView_0.1-9
##  [76] ProtGenerics_1.32.0     clue_0.3-65             leaps_3.1
##  [79] knitr_1.44              rstudioapi_0.15.0       tzdb_0.4.0
##  [82] coda_0.19-4             checkmate_2.2.0         nlme_3.1-163
##  [85] proxy_0.4-27            zoo_1.8-12              cachem_1.0.8
##  [88] parallel_4.3.1          miniUI_0.1.1.1          foreign_0.8-85
##  [91] vsn_3.68.0              pillar_1.9.0            grid_4.3.1
##  [94] vctrs_0.6.3             ggpubr_0.6.0            promises_1.2.1
##  [97] car_3.1-2               xtable_1.8-4            cluster_2.1.4
## [100] htmlTable_2.4.1         evaluate_0.21           mvtnorm_1.2-3
## [103] cli_3.6.1               compiler_4.3.1          rlang_1.1.1
## [106] crayon_1.5.2            ggsignif_0.6.4          labeling_0.4.3
## [109] affy_1.78.2             stringi_1.7.12          munsell_0.5.0
## [112] lazyeval_0.2.2          Matrix_1.6-1            hms_1.1.3
```

```
## [115] bit64_4.0.5            statmod_1.5.0          shiny_1.7.5
## [118] haven_2.5.3            broom_1.0.5            igraph_1.5.1
## [121] memoise_2.0.1          affyio_1.70.0          bit_4.0.5
## [124] readxl_1.4.3           ape_5.7-1
```
